# Marine biotoxin depuration rates: management applications, research priorities, and predictions for unstudied species

**DOI:** 10.64898/2025.12.16.694770

**Authors:** Christopher M. Free, Yutian Fang

## Abstract

Monitoring and managing the public health risk posed by marine biotoxins is a daunting challenge. With expansive coastlines, 1000s of seafood species, and dozens of biotoxins, monitoring cannot occur everywhere at once. Improving the cost-effectiveness of biotoxin monitoring is thus critical to expanding coverage to include more species, toxins, and locations. Notably, understanding the rate at which seafood species depurate biotoxins can be used to optimize the cadence of monitoring. We conducted a systematic review to collate marine biotoxin depuration rates and synthesize the factors that influence them; identify knowledge gaps, best practices, and research priorities; and generate predictions of depuration rates for unstudied species. We found that only 85 marine species have been studied for their biotoxin depuration rates. Depuration rates for non-bivalves and for toxins besides paralytic shellfish toxins (PSTs) are especially understudied. Depuration half-lives varied from 0.03 to 1269 days based on species, toxin, tissue, and environmental conditions. In general, depuration accelerates with increased temperature and food availability, with implications for aquaculture siting, depuration enhancement, and biotoxin monitoring. We identified unstudied bivalve and finfish species that are highly produced and vulnerable to HABs that are high priorities for future research. Finally, we used a Bayesian regression model to predict PST depuration rates for 102 unstudied bivalve species, the only clade-syndrome with sufficient training data. These predictions can guide efficient monitoring and management until lab- or field-based depuration rates become available. We recommend that future studies directly estimate depuration rates to ensure their comparability across studies and utility to managers.

## 1. Introduction

Harmful algal blooms (HABs) that produce biotoxins that accumulate in seafood species represent a growing threat to public health and marine fisheries and aquaculture (Hallegraeff et al., 2021). More than 200 taxa of marine algae produce biotoxins that threaten human health (Lundholm et al., 2009), with risk present in nearly all coastal waters (OBIS, 2025). Collectively, these toxins cause seven major poisoning syndromes in humans – paralytic, amnesic, diarrhetic, neurotoxic, azaspiracid, cyanotoxin, and ciguatera poisoning – with new toxins and syndromes discovered each decade (Nicolas et al., 2017). These syndromes cause symptoms ranging from gastrointestinal issues to neurological issues to death (Grattan et al., 2016). As a result, managers often close fisheries and aquaculture to protect public health, which can cause significant socioeconomic impacts from lost commercial, recreational, and cultural activities (Moore et al., 2020). HABs are expected to increase in frequency, duration, and intensity with climate change (IPCC, 2019), making them critical to study to ensure that seafood can safely remain a source of healthy, sustainable food for a growing human population.

To protect public health, many governments use biotoxin monitoring programs to track seafood safety and manage fisheries and aquaculture operations that are vulnerable to biotoxin risk (Andersen et al., 2004). In general, these programs monitor the toxicity of seafood species and trigger management actions when toxicity rises above a threshold deemed unsafe for human consumption, commonly referred to as the “action threshold” (Langlois and Morton, 2018). The management action, which could be a closure or an evisceration requirement (i.e., required removal of toxic body parts), is not ceased until toxicity falls below the action threshold, often after several successive tests to ensure safety. Ideally, all vulnerable species would be monitored to fully protect public health (Costa et al., 2017) and sampling would be highly spatially and temporally resolved to protect public health while also avoiding unnecessary management restrictions (Free et al., 2022). However, given limited resources to support expensive lab testing, expansive taxonomic, spatial, and temporal coverage is rarely possible (Langlois and Morton, 2018). Improving the cost-effectiveness of biotoxin monitoring is thus necessary to monitor more species, in more places, at more times. This could be achieved by either lowering the cost of each test (i.e., cheaper monitoring) or by lowering the number of tests required to effectively manage risk (i.e., more efficient monitoring).

One pathway for improving the efficiency of biotoxin monitoring programs is to tailor the frequency of sampling to the toxicokinetics of the monitored species (Blanco, 2009). Toxicokinetics describe the processes and rates of accumulation (a.k.a., uptake, adsorption), distribution, metabolism, and depuration (a.k.a., excretion, elimination, detoxification) of biotoxins within a species (Landrum et al., 1992). The rate at which species accumulate toxins determines how quickly monitoring should begin and proceed after a HAB is detected. For example, monitoring could begin later and advance more slowly if accumulation is slow but should begin sooner and advance more quickly if accumulation is fast. The distribution of toxin burden among different tissues is important because it determines what tissue is the most useful indicator of public health risk and whether a species could be safely consumed and successfully marketed following the evisceration (removal) of the most toxic tissue. The metabolism of the consumed toxin into new compounds, which can be either more or less toxic than the original compound (Bricelj and Shumway, 1998), contributes to the duration of seafood toxicity. Finally, the rate of depuration determines how frequently monitoring is needed after a species becomes toxic to facilitate the timely end of management actions. It can also be used to provide seafood harvesters, processors, and dealers forecasts of when management actions are likely to cease.

The use of depuration rates to design more efficient biotoxin monitoring and management is particularly promising because of the availability of good mathematical models for describing depuration and forecasting the duration of high toxicity. In general, depuration kinetics are theorized to follow either one- or two-compartment exponential decay (Blanco, 2009). In a one-compartment model, biotoxins are assumed to accumulate within a single “compartment”, which can represent a single tissue, a group of tissues, or even a whole organism, and are assumed to be exponentially lost to the environment at a single rate. For example, a toxin might accumulate in the digestive system following ingestion and then be quickly eliminated through egestion. In a two-compartment model, biotoxins are assumed to accumulate in an initial compartment (e.g., the digestive system) with some transference to a second compartment (e.g., muscle) with a slower exponential depuration rate. Theoretically, depuration rates could be described using even more compartments, though statistical support for increasingly complex models to offer more parsimonious fits seems unlikely; to our knowledge, the maximum number of compartments used in practice is three (Ye et al., 2021). With knowledge of the exponential decay constants for any of these models, managers could forecast depuration timelines (**Fig. 1**) and provide themselves, fishermen, and aquaculture farmers with valuable predictions of when seafood is expected to be safe for consumption.

**Figure 1.**
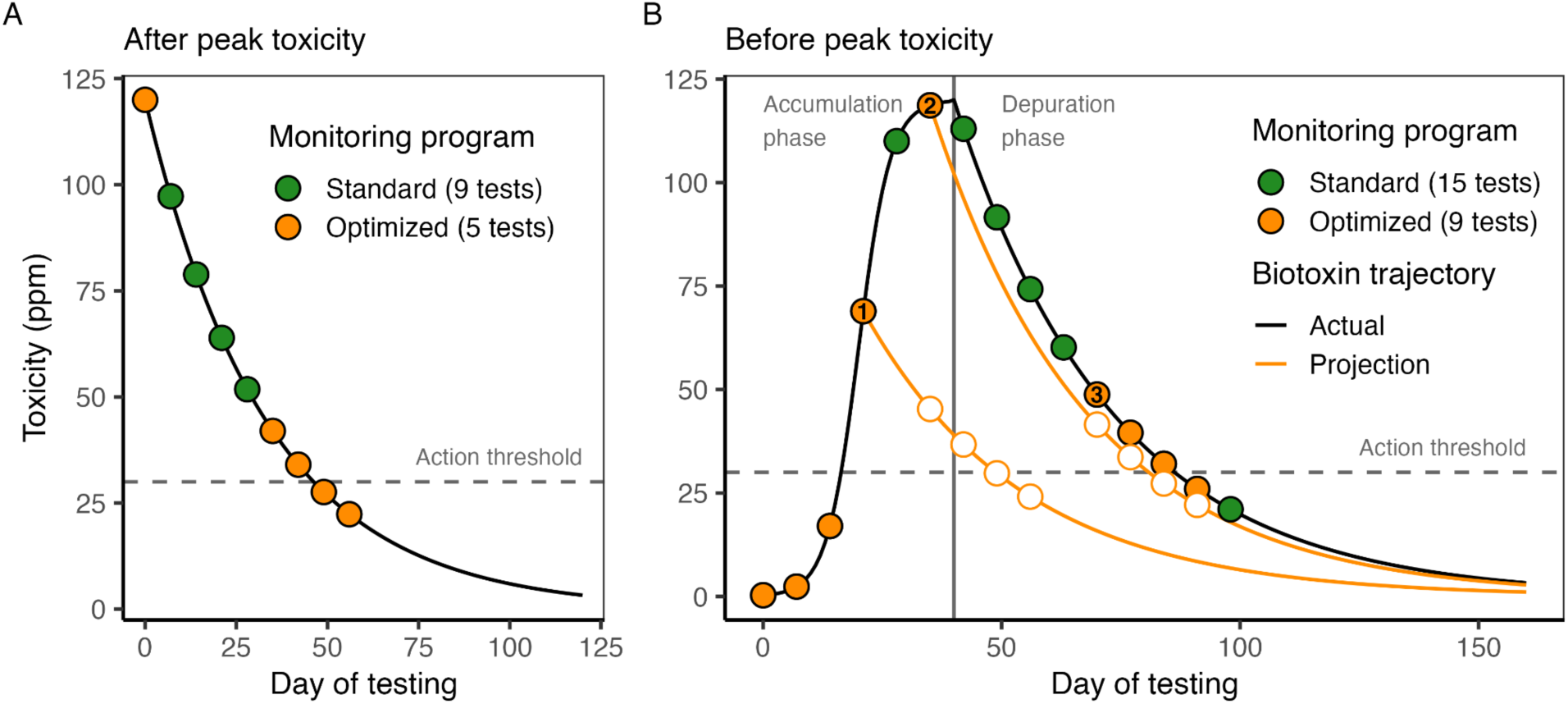
An illustration of the value of biotoxin depuration rates in efficiently monitoring and managing biotoxin risk both **(A)** after and **(B)** before peak toxicity is attained. In both panels, the black curve shows a hypothetical accumulation (B only) and depuration trajectory for a marine seafood species that depurates at 0.03 day^-1^. The points indicate weekly sampling that ceases once toxicity falls below the management action threshold (horizontal dashed line) in two consecutive weeks, a common requirement for opening a fishery closed due to biotoxin risk. In **(A)**, standard weekly sampling would require nine tests (all points), but pausing testing until two weeks before toxicity is projected to fall below the action threshold would require only five tests (orange points only). In **(B)**, scheduling tests based on three projections (marked by numbers) eliminates the need for six tests. The white circles indicate the projected testing schedule that is updated when toxicity is found to be higher than expected.

To illustrate how knowledge of depuration rates could improve the efficiency of biotoxin monitoring, we consider a species that depurates a biotoxin at 0.03 day^-1^ (2.9% day^-1^; 23.1 day half-life) based on one-compartment depuration kinetics (**Fig. 1**). This is roughly equivalent to the rate at which Atlantic surfclams (*Spisula solidissima*) depurate paralytic shellfish toxins (PSTs) under warm conditions (Bricelj et al., 2014). After reaching peak toxicity, managers could use this depuration rate to forecast when toxicity is expected to fall below the action threshold and tailor the cadence of monitoring accordingly (**Fig. 1A**). Specifically, managers could pause regular testing while toxicity is expected to remain well above the action threshold and resume it when toxicity is expected to approach the threshold. In this example, optimized testing eliminates the need for four tests, which accumulates when applied across many management zones (e.g., 10 zones = 40 saved tests). This frees up resources to monitor new management zones or new species at the same cost as the original monitoring program. Importantly, knowledge of depuration rates can improve efficiency even before peak toxicity is attained (**Fig. 1B**). If the first test conducted after a forecast is higher than expected (because the species is still accumulating toxins from an ongoing bloom or from toxins in the food web), the forecast can be repeated and the cadence of monitoring updated accordingly.

The widespread use of depuration rates to increase the efficiency of biotoxin monitoring programs requires the consolidation of known depuration rates, identification of the factors that affect biotoxin depuration, and prioritization of depuration rates still needing study. This is partially supported by a small number of review papers. A review by Fernandez et al. (2003) provided biotoxin “retention times”, the time required for toxicity to fall below either the action threshold or the limit of detection, for 34 species-toxin combinations representing 31 bivalve species and five toxin syndromes (paralytic, diarrhetic, amnesic, neurotoxic, other). Although these values provide credible evidence that depuration rates are species- and toxin-specific, they are challenging to interpret and cannot be operationalized as rates. A review by Bricelj and Shumway (1998) provides operational PST depuration rates (% per day) for 21 species of bivalves. A handful of research papers provide non-systematically collated depuration rates to place measured depuration rates within a broader context (Garcia-Corona et al., 2024; Martins et al., 2020; Schultz et al., 2013). While valuable, these reviews collectively lack depuration rates for non-bivalves, three notable biotoxin syndromes (ciguatera, cyanotoxin, azaspiracid), and over two decades of research into marine biotoxin depuration processes.

Here, we seek to empower more efficient marine biotoxin monitoring and management through a multi-pronged assessment of biotoxin depuration rates. First, we conducted a systematic literature review to collate published depuration rates, synthesize information on the factors that impact biotoxin depuration, and identify best practices for future depuration studies. Second, we prioritized species for future depuration study by identifying highly produced seafood species that are vulnerable and exposed to each biotoxin syndrome yet have no published depuration rates. Finally, we used a Bayesian regression model trained on our database of biotoxin depuration rates to predict rates for unstudied species, which can be used to guide monitoring programs until depuration studies are completed. Overall, we demonstrate how biotoxin depuration rates can forecast depuration timelines, which can be used to design more efficient monitoring programs and to support informed management and business decisions.

## 2. Methods

### 2.1 Literature review

#### 2.1.1 Database development

We conducted a systematic review of marine biotoxin depuration rates following the PRISMA review protocol (**Fig. S1**) (Page et al., 2021). First, we identified 797 candidate papers by using the following search query in Web of Science on July 30, 2025: (“depurat*” or “excret*” or “eliminat*” or “detox*”) AND (“toxin*” or “biotoxin*” or “phycotoxin” or “domoic” or “okadaic” or “saxitoxin” or “brevetoxin” or “azaspiracid” or “cyanotoxin” or “ciguatoxin”) AND (“marine” or “ocean”). This query was designed to identify papers that study the depuration (a.k.a. excretion, elimination, or detoxification; term 1) of marine phycotoxins (term 2) from marine species (term 3). We only considered papers that either quantified the depuration rates of biotoxins produced by harmful algae from marine species or presented the data required for us to externally quantify depuration rates. Thus, we excluded 643 papers that did not study a marine species, did not study a biotoxin produced by a harmful algae species, or did not quantify a depuration rate or present the data needed to externally derive a depuration rate (**Fig. S1**). We supplemented this systematic review with a targeted search of the Chinese literature to identify depuration rates not represented in the English literature; this non-systematic review added two papers on paralytic shellfish toxin depuration rates in Manila clam (*Ruditapes philippinarum*).

We reviewed the resulting 156 papers (**Fig. S2**) and extracted the following attributes of the study (**Table S1**): species studied (common and scientific name), biotoxin studied and its causative agent, tissue studied (e.g., meat, viscera, hepatopancreas; **Table S2**), study type (lab or field), feeding conditions during depuration (starved, fed a non-toxic diet, or foraged in the wild), and other experimental conditions (e.g., varied temperature, salinity, diets, etc.). If the depuration rate (exponential decay constant or half-life) was directly reported, we recorded the rate and whether it was derived from a one- or two-compartment exponential decay model (details below). These are also known as first- or second-order or one- or two-phase (biphasic) decay models. We grouped the studied biotoxins into eight syndromes of shellfish poisoning (Nicolas et al., 2017): amnesic, diarrhetic, paralytic, ciguatera, neurotoxic, azaspiracid, cyanotoxin, and other shellfish poisoning (**Table 1**). Cyanotoxin poisoning describes toxins formed by cyano-bacteria including microcystin, nodularin, and homoanatoxin. The other category includes yessotoxins, pectenotoxins, tetrodotoxins, gymnodimines, and karlotoxins.

**Table 1.** Marine biotoxin poisoning syndromes and their causative organisms.

| Syndrome | Biotoxin | Causative organisms |
| --- | --- | --- |
| Amnesic (ASP) | Domoic acid (DA) | Diatom: *Pseudo-nitzschia* spp. |
| Diarrhetic (DSP) | Diarrhetic shellfish toxins (DSTs) | Dinoflagellate: *Dinophysis* spp., *Prorocentrum* spp. |
| Paralytic (PSP) | Paralytic shellfish toxins (PSTs)<sup>2</sup> | Dinoflagellate: *Alexandrium* spp, *Pyrodinium bahamense*, *Gymnodinium catenatum* |
| Azaspiracid (AZP) | Azaspiracid (AZA) | Dinoflagellate: *Azadinium* spp., *Amphidoma spinosum* |
| Ciguatera (CFP) | Ciguatoxin (CTX) | Dinoflagellate: *Gambierdiscus* spp., *Fukuyoa* spp. |
| Neurotoxic (NSP) | Brevetoxin (PbTx) | Dinoflagellate: *Karenia* spp. |
| Cyanotoxin | Cyanotoxins<sup>3</sup> | Cyanobacteria |
| Other | Other phycotoxins<sup>4</sup> | Dinoflagellate: *Protoceratium reticulatum*, *Lingulodinium polyedra*, *Gonyaulax* spp. |

<sup>1</sup> Okadaic acid (OA), dinophysistoxin (DTX)
<sup>2</sup> Saxitoxin (STX), gonyautoxin (GTX)
<sup>3</sup> Microcystin, nodularin, homoanatoxin, cylindrospermopsin
<sup>4</sup> Yessotoxin (YTX), pectenotoxin (PTX, associated with DSTs but do not cause DSP), tetrodotoxin (TTX), gymnodimine (GYM), karlotoxin

#### 2.1.2 Deriving depuration rates

If the depuration rate was not directly reported as either an exponential decay constant or half-life, we extracted the data needed to derive the depuration rate using WebPlotDigitizer (Rohatgi, 2025). We estimated depuration rates using a one-compartment exponential decay model fit to the depuration phase of the data. For field studies with long time series, depuration rates were often estimated for discrete depuration events **(Fig. S3**). We calculated tissue-specific depuration rates such that:

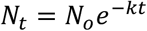

where toxicity at time *t* (*N_t_*) is a function of initial toxicity (*N_0_*) and the decay constant *k*.

We calculated the half-life, the time required for toxicity to halve, associated with each reported or derived one-compartment model depuration rate as:

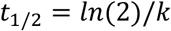

If a paper reported a half-life but not the decay constant (*k*), we inverted this equation to calculate the decay constant. We standardized all decay constants and half-lives to be in terms of days (i.e., some fast depuration rates were reported or derived in terms of hours). Finally, we calculated the percent daily loss of biotoxin burden as:

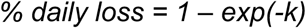

### 2.2 Priority species for depuration study

We identified priority species for depuration study as vulnerable species that have high commercial harvests from countries with HAB exposure but that lack published depuration rates. First, we identified countries exposed to HAB species associated with each biotoxin syndrome (**Table 1**) using data from the Ocean Biodiversity Information System (OBIS) (OBIS, 2025), which provides geotagged observations of HAB species (**Fig. S4**). We classified a country’s waters as exposed to biotoxin syndrome if an associated species (OBIS) has ever been recorded within its Exclusive Economic Zone (EEZ) (Flanders Marine Institute, 2025). We note that this likely underestimates countries exposed to HABs as many countries, especially in Africa and the Middle East **(Fig. S4**), lack HAB monitoring programs (Andersen et al., 2004).

Second, we identified shellfish and finfish species that are vulnerable to HABs and harvested in exposed countries using national fisheries and aquaculture production data from the Food and Agriculture Organization (FAO) of the United Nations (FAO, 2024). We identified vulnerable shellfish as filter-feeding molluscs (e.g., oysters, mussels, clams, scallops), as they readily accumulate toxins by directly consuming HAB species, even within aquaculture facilities where they generally feed directly from the surrounding environment. We excluded predatory molluscs (e.g., cephalopods), grazing molluscs (e.g., gastropods), and non-mollusc shellfish (e.g., crustaceans, echinoderms), which are challenging to prioritize because their indirect biotoxin accumulation pathways make their vulnerability to biotoxins highly heterogeneous. The shellfish prioritization excludes ciguatera, which is principally a finfish biotoxin syndrome.

We identified vulnerable finfish as those vulnerable to ciguatera, the primary biotoxin syndrome affecting finfish. Ciguatera is most common in large, tropical, reef-associated, predators (Lewis and Holmes, 1993; Randall, 1958). We thus used information on habitat associations, maximum length, and trophic level from FishBase (Boettiger et al., 2012; Froese and Pauly, 2025) to identify reef-associated predators larger than 25 cm that are harvested in capture fisheries. A threshold of 25 cm was selected as the smallest fish with detectable ciguatoxins in the review of (Li et al., 2023) was striated surgeonfish (*Ctenochaetus striatus*) at 26 cm maximum length. Because FishBase information on trophic level is relatively sparse, we could only exclude known herbivores (i.e., we could not filter for known predators). FishBase also identifies fish with known ciguatera observations; these species were considered in the prioritization even if they did not meet the criteria above. We excluded aquaculture production from the finfish prioritization analysis because predatory finfish are generally fed non-toxic food and are thus less exposed to ciguatoxins through the food web.

Finally, we identified the 20 vulnerable species with the highest annual production (bivalves: aquaculture+fisheries; finfish: fisheries only) from 2014-2023 in EEZs exposed to each biotoxin syndrome and classified species without published depuration rates as priorities for study. We highlighted unstudied species that would provide the first depuration estimate at the order, family, or genus levels as particularly valuable. To illustrate the value of these estimates, we quantified the (1) number of currently harvested species in the taxonomic group using the FAO data and (2) total number of species in the group using FishBase and SeaLifeBase (Palomares and Pauly, 2025), which captures potential future harvest diversity.

Although we did not prioritize non-bivalve invertebrates for depuration study, we compared maximum biotoxin toxicities reported in non-bivalves by select review papers (Costa et al., 2017; Deeds et al., 2008; Lefebvre and Robertson, 2010) relative to common international action thresholds (Langlois and Morton, 2018) to assess the importance of conducting depuration studies for non-bivalve vectors of seafood poisoning syndromes. These review papers were opportunistically selected from the systematic literature search but do not constitute a systematic review of all observed non-bivalve toxicities.

### 2.3 Predicting bivalve PST depuration rates

We compared the ability for one generalized linear model and three mixed effects models to predict paralytic shellfish toxin (PST) depuration rates in marine bivalves. We focused on PST depuration rates in bivalves based on the *a priori* expectation that this was the only clade-syndrome combination with sufficient data (**Fig. 2**) to generate a model with good predictive skill. All four models predict the log depuration rate to normalize the response variable and to constrain predictions to positive depuration rates (i.e., exponential decay rather than exponential growth). All four models include five fixed effects: (1) study type (lab vs. field), (2) tissue type (e.g. muscle, hepatopancreas, etc.); (3) maximum length (cm), (4) von Bertalanffy somatic growth rate (1/yr), which describes the rate at which an organism grows towards its maximum size, and (5) preferred temperature (°C). These variables were selected based on hypotheses that (a) depuration rates would be faster in the lab, where species are fed non-toxic diets; (b) depuration rates would vary by tissue; and (c) depuration rates would be faster for smaller, faster growing, warmer water species, which are expected to have faster metabolisms (Robinson et al., 1983) and therefore faster egestion. We retrieved the maximum length, growth rate, and preferred temperature from SeaLifeBase (Palomares and Pauly, 2025) using the *rfishbase* R package (Boettiger et al., 2012). When values were missing for a species, we preferentially used the genus-, family-, order-, or class-level median, depending on availability.

**Figure 2.**
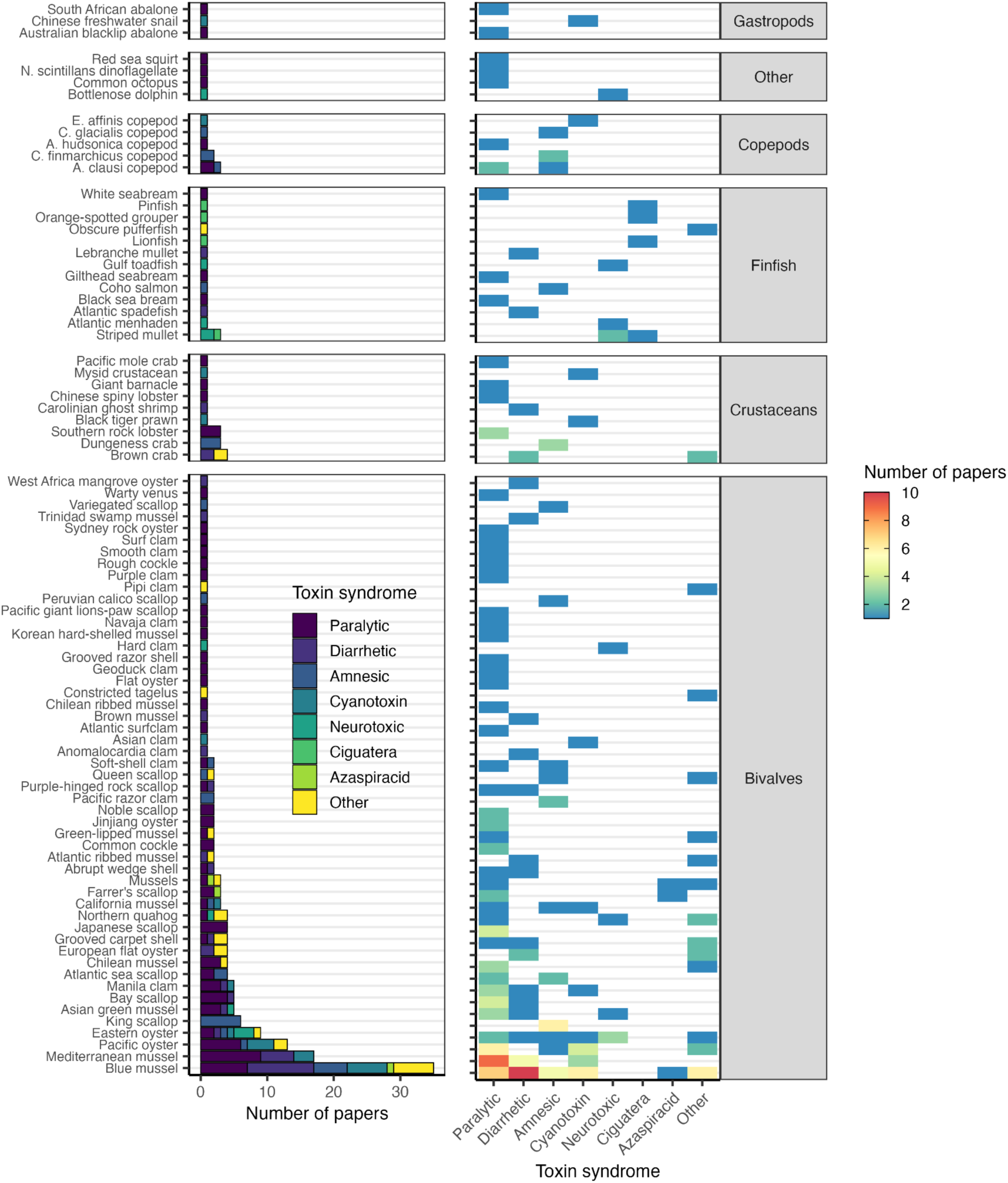
The number of papers measuring depuration rates by species and biotoxin. Species are grouped by taxonomic class and are ordered by increasing sample size (bottom to top). Biotoxins are also ordered by increasing sample size (left to right). See **Table S3** for the scientific names of all species shown here.

The first model includes only the five fixed effects variables (the “fixed effects only” model) while the subsequent models use different approaches for incorporating taxonomic random effects. The inclusion of taxonomic random effects evaluates the hypothesis that traits besides those explicitly evaluated as fixed effects may also determine depuration rates. For example, species-specific physiology and feeding ecology may play an important role in depuration, but we lack the data to explicitly incorporate these traits as fixed effects. The “phylogenetic random effects” model evaluates the hypothesis that depuration rates are phylogenetically conserved and correlated to ancestry. This model includes species-specific random effects that were modelled using a variance-covariance matrix derived from the bivalve phylogenetic tree. We constructed a phylogenetic tree of all bivalve species using the Open Tree of Life API (Hinchliff et al., 2015) via the *rotl* R package (Michonneau et al., 2016). We estimated branch lengths using the Grafen method (Grafen, 1997), which scales internal nodes to approximate evolutionary divergence when branch-length information is incomplete, implemented through the *ape* R package (Paradis and Schliep, 2019). The “taxonomically nested model” has hierarchically nested order, family, genus, and species random effects. This model evaluates the hypothesis that depuration rates are similar within a clade but that differences between clades are not related to ancestry (as in the phylogenetic model). Finally, the “species random effects” model includes species-level random effects that are not structured by phylogeny or taxonomy. This model evaluates the hypothesis that unobserved species traits impact depuration rates but that these impacts are idiosyncratic and not related to taxonomy.

We fit all four models using a Bayesian estimation approach implemented through the *brms* R package (Bürkner, 2017). The models were compared using efficient approximate leave-one-out cross-validation (LOO) using the *loo* R package (Vehtari et al., 2024), where the best model was identified as the one with the largest expected log predictive density (ELPD), which characterizes the ability of the model to predict the data withheld in the cross-validation. The fit of all four models was also evaluated through the estimation of a Bayesian R^2^ (Gelman et al., 2019). We evaluated the coefficients and conditional effects of predictors in the best fitting model to understand the impact of each variable on PST depuration rates. Finally, we used the best fitting model to estimate PST depuration rates for all harvested marine bivalves, as determined through the analysis of the FAO production data (FAO, 2024) described above. We included all harvested bivalves, not just those harvested from EEZs with known PST occurrence (**Fig. S4**), given that PST may be present but undetected in many of these EEZs and the potential for the range of PST causative agents to expand in the future. The life history traits for these species were also retrieved from SeaLifeBase using the *rfishbase* R package.

## 3. Results

### 3.1 Literature review

#### 3.1.1 General characteristics

Biotoxin depuration rates have been studied in 85 marine species spanning 66 genera, 39 families, 26 orders, and 10 classes (**Fig. 2A; Table S3**). Marine invertebrates, especially bivalves, have been much more studied than marine vertebrates: only 13 papers (8%) have evaluated biotoxin depuration rates in finfish whereas 145 papers (93%) have assessed biotoxin depuration rates in invertebrates (some papers studied both an invertebrate and a vertebrate allowing this statistic to sum to >100%; this applies to other results as well). Most of the evaluated papers studied bivalves (124 papers; 80%), with a particular focus on blue mussels (31 papers; 20%), Mediterranean mussels (17 papers; 11%), Pacific oysters (13 papers; 8%), eastern oysters (8 papers; 5%), and king scallops (6 papers; 4%). Notably, blue mussels are the only species with depuration rates estimated for six of the eight biotoxin syndromes (**Fig. 2B**).

The depuration of PSTs from marine species has been studied more than the depuration of any other biotoxin (66 papers; 42%) (**Fig. 2B**). Domoic acid (24 papers, 15%), diarrhetic shellfish toxin (DST) (24 papers; 15%), and cyanotoxin (16 papers; 10%) depuration rates have received similar levels of attention. Brevetoxin depuration has been evaluated in 9 papers (6%), which evaluate four bivalve, two finfish, and one dolphin species. Ciguatoxin depuration has only been studied in four finfish species (4 papers) and azaspiracid depuration has only been measured in three bivalve species (3 papers). Eighteen papers (12%) have assessed the depuration rates of “other” biotoxins (**Table 1**), which include yessotoxins, tetrodotoxins, pectenotoxins, karlotoxins, and gymnodimines (**Fig. 2B**).

Biotoxin depuration has been studied in many tissue types (**Fig. S5**) with some papers comparing depuration rates among many tissues (**Fig. 3B**). In general, the tissues most evaluated are those that either harbor the greatest toxin burden or are consumed by people. For bivalves, studies have largely focused on the soft tissue (eaten by people) and the hepatopancreas (greatest toxin burden; Bricelj and Shumway, 1998). Similarly, crustacean depuration rates have largely been evaluated in the hepatopancreas (greatest toxin burden; Schultz et al., 2013) and the soft tissue (eaten by people). For finfish, the dominant tissues studied are the liver and muscle (eaten by people). Most gastropod studies have measured depuration in the foot (eaten by people) or the whole organism. All cephalopod studies have focused exclusively on the viscera, and all phytoplankton, zooplankton, and sea squirt studies have exclusively analysed the whole organism (**Fig. S6**).

**Figure 3.**
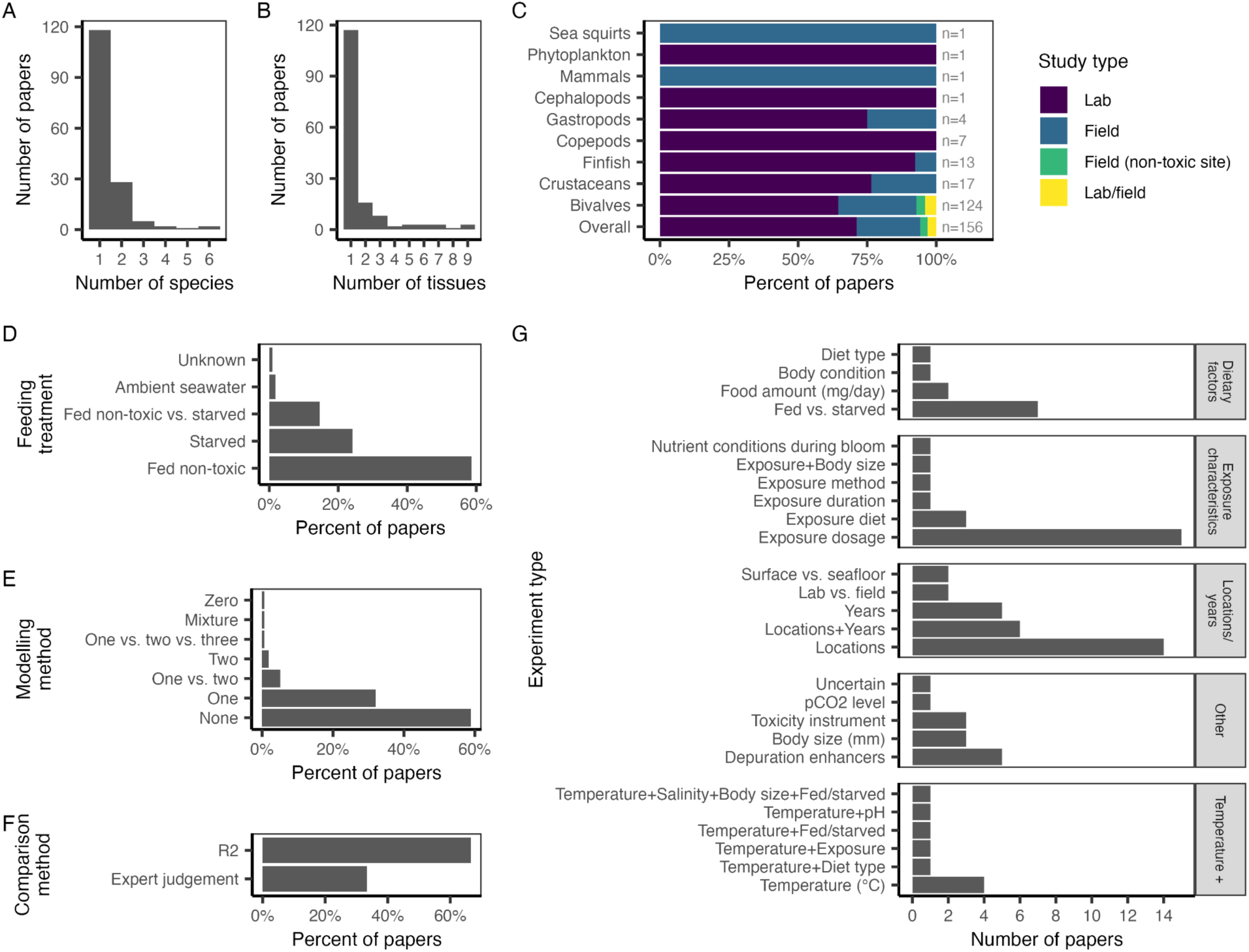
Characteristics of papers measuring biotoxin depuration in marine species. Panel **A** shows the number of species studied in a paper. Panel **B** shows the number of tissues evaluated in a paper. Panel **C** shows the percent of papers that conduct lab and/or field studies of depuration rates; a few studies conduct depuration studies at non-toxic field sites. Panel **D** shows the feeding treatment used during the depuration phase of each study; all field studies use wild diets. Panel **E** shows the modeling method used to quantify depuration rates (i.e., one-or two-compartment exponential decay models). Panel **F** shows the method used to compare models with different numbers of compartments in papers making such comparisons. Panel **G** shows the number of studies conducting experiments to measure depuration rates under different conditions. Experiment types are grouped into broader categories.

Most depuration studies occur in the lab (71%; 110 papers), where depuration can be measured under controlled diet and environmental conditions (**Fig. 3C**). 23% (36 papers) of depuration studies occur in the field, where organisms are potentially exposed to toxic algae while depurating (i.e., depuration may be offset by uptake), but where depuration is quantified in a real-world setting. Four studies (3%) depurate organisms fed toxic algae in the lab at non-toxic field sites to mimic depuration in a natural, but non-toxic, setting. Five studies (3%) directly compared lab and field depuration rates. Houle et al. (2023) find that PST depuration in Purple-hinged rock scallop (*Crassadoma gigantea*) is 3.1 times slower in the field. Slower depuration in the field is also found for the depuration of domoic acid by Peruvian calico scallops (*Argopecten purpuratus*) (Alvarez et al., 2020) and variegated scallops (*Mimachlamys varia*) (Le Moan et al., 2025) and the depuration of PSTs from purple clams (*Hiatula rostrata*) (Chen and Chou, 2002) and blue mussel (*Mytilus edulis*) (Scarratt et al., 1991). Our meta-analysis confirms this result for most species with depuration rates estimated for common tissues in field and lab studies (**Fig. S6**).

#### 3.1.2 Experimental results

Many of the evaluated papers assessed depuration rates under different experimental treatments (**Fig. 3G**). In particular, many lab studies considered the impacts of water temperature, dietary factors, and biotoxin exposure characteristics on depuration rates. A few lab studies also considered the impacts of size and/or age on depuration rates, the ability for industrial additives to enhance depuration rates, and the sensitivity of depuration rate estimates to measurement technique. Many field studies assessed depuration rates across different locations and/or years and two field studies compared depuration rates at the seafloor and surface. Five studies directly compared depuration rates in the lab, where organisms are fed a controlled non-toxic diet, versus the field, where organisms experience a natural diet that may not be completely toxin-free (**Fig. 3G**).

##### 3.1.2.1 Impacts of exposure intensity

Because studies of biotoxin depuration rates are often conducted following studies of biotoxin accumulation (e.g. Kwong et al., 2006; Lopes et al., 2014a; Lund et al., 1997), many of the evaluated studies (n=22) included trials with different exposure intensities. However, the primary purpose of these treatments is to quantify feeding clearance rates and associated biotoxin uptake rates, which are density-dependent. Although these treatments impact the peak toxicity (*N0*) of a one-compartment depuration model, they would not be expected to affect the depuration rate constant (*k*), which is, by definition, the same at any toxicity. As a result, most of these papers do not explicitly compare depuration rates across treatments and we made no attempt to make these comparisons ourselves. Most of these studies (n=16) manipulated exposure intensity by varying the density (dosage) of the causative agent. The remaining studies varied exposure intensity by varying the composition of causative agents in the diet (n=3; Bricelj et al., 1991; Jauffrais et al., 2012; Lewis et al., 2022), the duration of exposure to the causative agent (n=1; Lin et al., 2024), the nutrient conditions during exposure (n=1; Strogyloudi et al., 2006), or the method of exposure (n=1; i.e., oral exposure vs. injection; Kankaanpää et al., 2005).

##### 3.1.2.2 Impacts of dietary factors

The eleven papers that evaluate the impact of dietary factors on biotoxin depuration rates generally find that individuals depurate faster when fed than when starved and that they depurate even faster when provided more food or when hungrier. This is generally believed to occur because depuration largely occurs through egestion of feces or pseudofeces and egestion is generally higher when eating (Bricelj and Shumway, 1998). Faster depuration has been found for fed versus starved individuals for: (i) PSTs in Pacific oysters (*Crassostrea gigas*) (Medhioub et al., 2012) and Jinjiang oysters (*Ostrea rivularis*) (Yang et al., 2021); (ii) okadaic acid in blue mussels (Marcaillou et al., 2010); (iii) domoic acid in Dungeness crab (*Metacarcinus magister*) (Lund et al., 1997); and (iv) gymnodimines in grooved carpet shells (*Ruditapes decussatus*) (Medhioub et al., 2010). Blue mussels depurated okadaic acid faster when fed more food (Marcaillou et al., 2010). Relatedly, field-collected brown crabs (*Cancer pagrus*) in poor body condition (hungrier) depurated faster okadaic acid faster than lab-fed brown crabs in better body condition (less hungry). The depuration rate of okadaic acid (Svensson, 2003) and domoic acid (Wohlgeschaffen et al., 1992) from blue mussels and of domoic acid from three *Calanus* copepod species (*C. finmarchicus, C. glacialis, C. finmarchicus*) (Hardardottir et al., 2019; Leandro et al., 2010) were all so rapid that they were not affected by whether they were starved or fed. Only northern quahog (*Mercenaria mercenaria*) and constricted tagelus (*Sinonovacula constricta*) were found to depurate karlotoxins faster when starved than when fed (Li et al., 2024). The authors hypothesize that this surprising finding may be because stomach contents, including residual toxic algae, are evacuated more efficiently when starved.

##### 3.1.2.3 Impacts of temperature

The nine papers that evaluated the impact of temperature on biotoxin rates generally find that depuration is faster in warmer waters. Blue mussels (*Mytilus edulis*) depurated domoic acid faster at 11°C than 6°C (Novaczek et al., 1992) and faster at 11°C than 6°C (Scarratt et al., 1991). Similarly, wedge shells (*Donax trunculus*) depurated okadaic acid 71% faster at 20°C than at 17°C (Botelho et al., 2018). Atlantic surfclams (*Spisula solidissima*) viscera, where toxicity is highest, depurated okadaic acid faster under warmer conditions though temperature did not impact depuration rates in other tissues (Bricelj et al., 2014). On the other hand, Svensson and Förlin (2004) found that even though blue mussels (*Mytilus edulis*) eliminated lipids faster in warmer waters, water temperature did not affect the depuration rate of okadaic acid, a lipophilic phycotoxin. Farrell et al. (2015) found that Sydney rock oyster (*Saccostrea glomerata*) and diploid Pacific oysters (*Crassostrea gigas*) depurated PST faster at 27°C than at 22°C, but that temperature did not affect depuration rates for triploid Pacific oysters. Tang et al. (2021b) found that Korean hard-shelled mussel (*Mytilus coruscus*) depurated PSTs faster at 30°C than at 25°C. In the only study examining the impact of temperature on the biotoxin depuration rate of a finfish, Barbosa et al. (2019) found that gilthead seabream (*Sparus aurata*) depurated PST so quickly that an impact of temperature could not be measured. Finally, Braga et al. (2018) is the only paper to find slower depuration under warming, where Mediterranean mussels depurated PSTs faster at 19°C than at 24°C.

##### 3.1.2.4 Impacts of body size and age

The five papers that considered the impact of size or age on biotoxin depuration rates generally found that smaller/younger individuals depurate faster than larger/older individuals. Bogan et al., (2007) found that smaller king scallops (*Pecten maximus*) depurated domoic acid faster than larger scallops. Mafra et al. (2010) found that smaller Eastern oysters (*Crassostrea virginica*) depurated domoic acid faster than larger oysters but that size did not affect the domoic acid depuration rates of blue mussels (*Mytilus edulis*). Similarly, Duinker et al. (2007a) found no difference in the okadaic acid depuration rates of 1- and 2-year-old blue mussels. Finally, Min et al. (2018) show that cyanotoxin depuration began earlier for juvenile mysid crustaceans (*Neomysis awatschensis*) and thus progressed faster.

##### 3.1.2.5 Impacts of depuration enhancers

A few papers examined the ability for additives to enhance the depuration process. Xie et al. (2013) found that supplementing Jinjiang oyster (*Ostrea rivularis*) diets with chitosan, a natural polymer that adsorbs PST toxins and prevents their reuptake by depurating filter feeders, significantly accelerated their PST depuration. Qiu et al. (2018) found that adding activated carbon, which also adsorbs PST toxins and prevents their reuptake by depurating filter feeders, to depuration tanks significantly accelerated PST depuration from both Mediterranean mussels (*Mytilus galloprovincialis*) and Farrer’s scallops (*Chlamys farreri*). Activated carbon is a highly effective biotoxin adsorbent because of its high porosity, surface area, and cationic charge. Similarly, Peña-Llopis et al. (2014) found that adding N-Acetylcysteine, which boosts the production of compounds central to the detoxification process, resulted in a four-fold increase in the depuration of domoic acid from king scallops (*Pecten maximus*), one of the slowest depurating bivalve species (Blanco et al., 2002). On the other hand, Leal et al. (2023) found that adding cation-exchange resins (CER), which adsorb positively charged toxic compounds, did not significantly accelerate PST depuration in blue mussel (*Mytilus edulis*).

#### 3.1.3 Depuration rates by species

Only 41% (n=64) of the evaluated studies reported biotoxin depuration rates in terms of exponential decay constants or half-lives, which are quantitative values that can be directly compared across studies, species, tissues, experimental conditions, biotoxins, etc. (**Fig. 3E**). The remaining 59% of studies (n=92) described depuration rates in bespoke terms that are not directly comparable across studies, such as “*after 14 days, the [domoic acid] concentration in the fed crabs decreased by 73%*” (Lund et al., 1997) or “*[toxicity decreased]* f*rom 3.1 μg [okadaic acid] g^-1^ …. at day 1 to 1.51 μg [okadaic acid] g^-1^ at day 32*” (Svensson, 2003). Among the 66 studies that reported exponential decay constants or half-lives, 50 (76%) estimated these values using one-compartment exponential decay models, 3 (5%) estimated these values using two-compartment models (Choi et al., 2003; Schultz et al., 2008; Yu et al., 2005), 8 (12%) compared the results of one- and two-compartment models (Alvarez et al., 2020; Blanco et al., 1999; Jauffrais et al., 2012; Lopes et al., 2014; Mafra et al., 2010; Moroño et al., 2003; Nielsen et al., 2016; Woofter et al., 2005), and one compared the results of one-, two-, and three-compartment models (Kennedy et al., 1992). Only four of the studies comparing one- and multi-compartment models found significantly more support for the multi-compartment model (Alvarez et al., 2020; Jauffrais et al., 2012; Kennedy et al., 1992; Woofter et al., 2005). These comparisons were made on a mixture of expert judgement and comparison of R^2^ values, which also involved extensive expert judgement (**Fig. 3F**). Note that many studies (n=39) compared depuration rates among different tissues (**Fig. 3B**), which also provides information on multi-compartment depuration rates.

Depuration rates varied from 22.3 day^-1^ (0.03 day half-life) for microcystin from whole *Eurytemora affinis* copepods (Karjalainen et al., 2006) to 0.0005 day^-1^ (1269 day half-life) for domoic acid from the gills of Dungeness crab (*Metacarcinus magister*) (Schultz et al., 2013) (**Fig. 4**). Depuration rates appeared relatively similar among species of the same genus (**Fig. 4**), motivating the taxonomic regression analysis described in *Section 2.3*.

**Figure 4.**
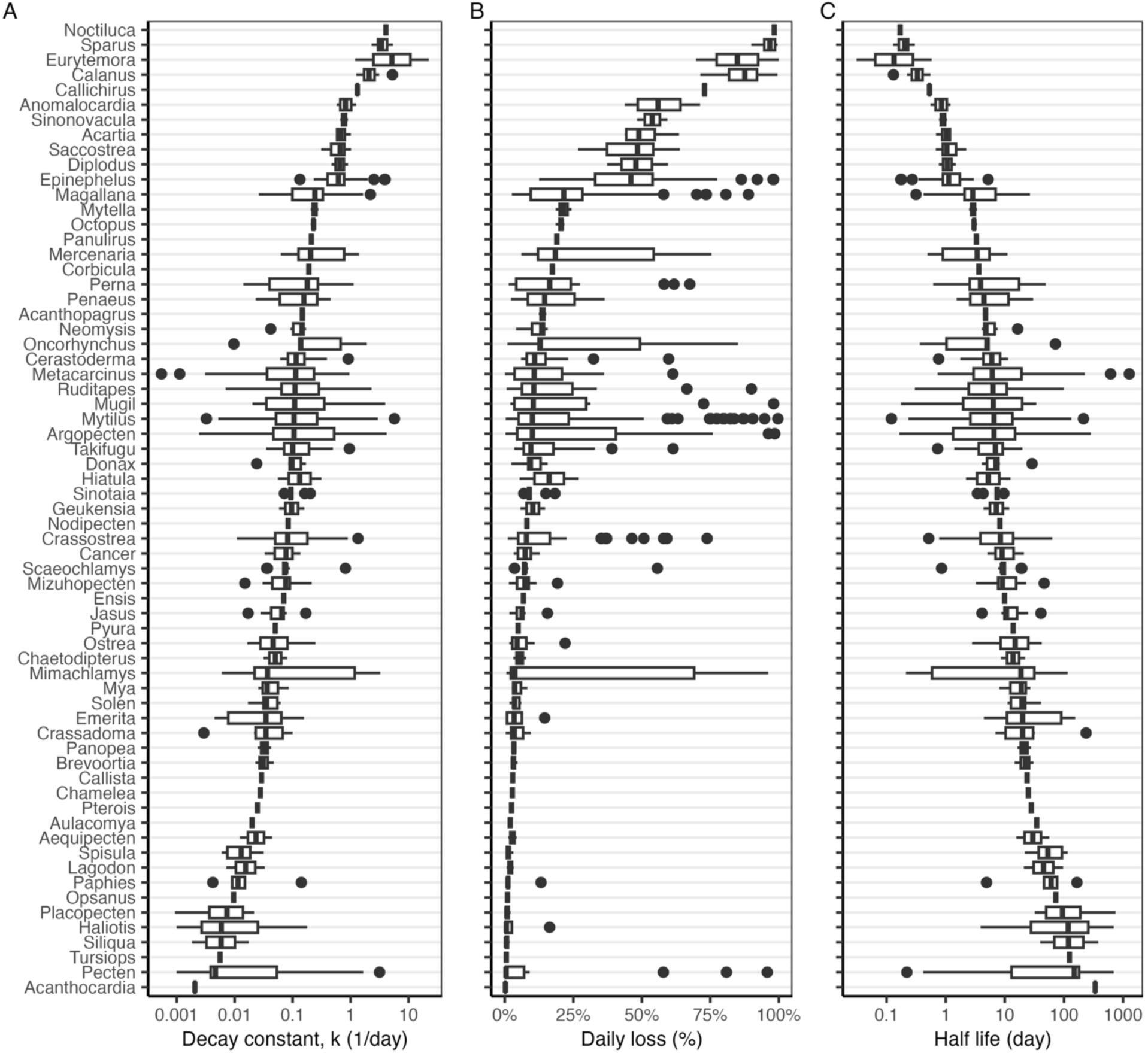
The **(A)** exponential decay constant (k, day^-1^), **(B)** daily percent loss (%), and **(C)** half-life (days) of marine biotoxins by genus. In the boxplots, the solid line indicates the median, the box indicates the interquartile range (IQR; 25th to 75th percentiles), the whiskers indicate 1.5 times the IQR, and points indicate outliers.

### 3.2 Priority species for depuration study

Thirteen of the top-20 most harvested marine filter-feeding mollusc species from countries exposed to PSTs have had their PST depuration rates studied (**Fig. 5A**), the highest of any of the biotoxin types (**Fig. 5**). However, three of the top-5 species have not had their PST depuration rates examined: constricted tagelus (*Sinonovacula constricta*) and blood cockle (*Tegillarca granosa*). PST depuration rates for 1 new order, 2 new families, and 4 new genera could be gained through the evaluation of 7 of the 10 unstudied species (**Table S4**). In particular, evaluation of blood cockle (*Tegillarca granosa*) would provide information for 8 harvested and 360 total species in the Arcoida order.

**Figure 5.**
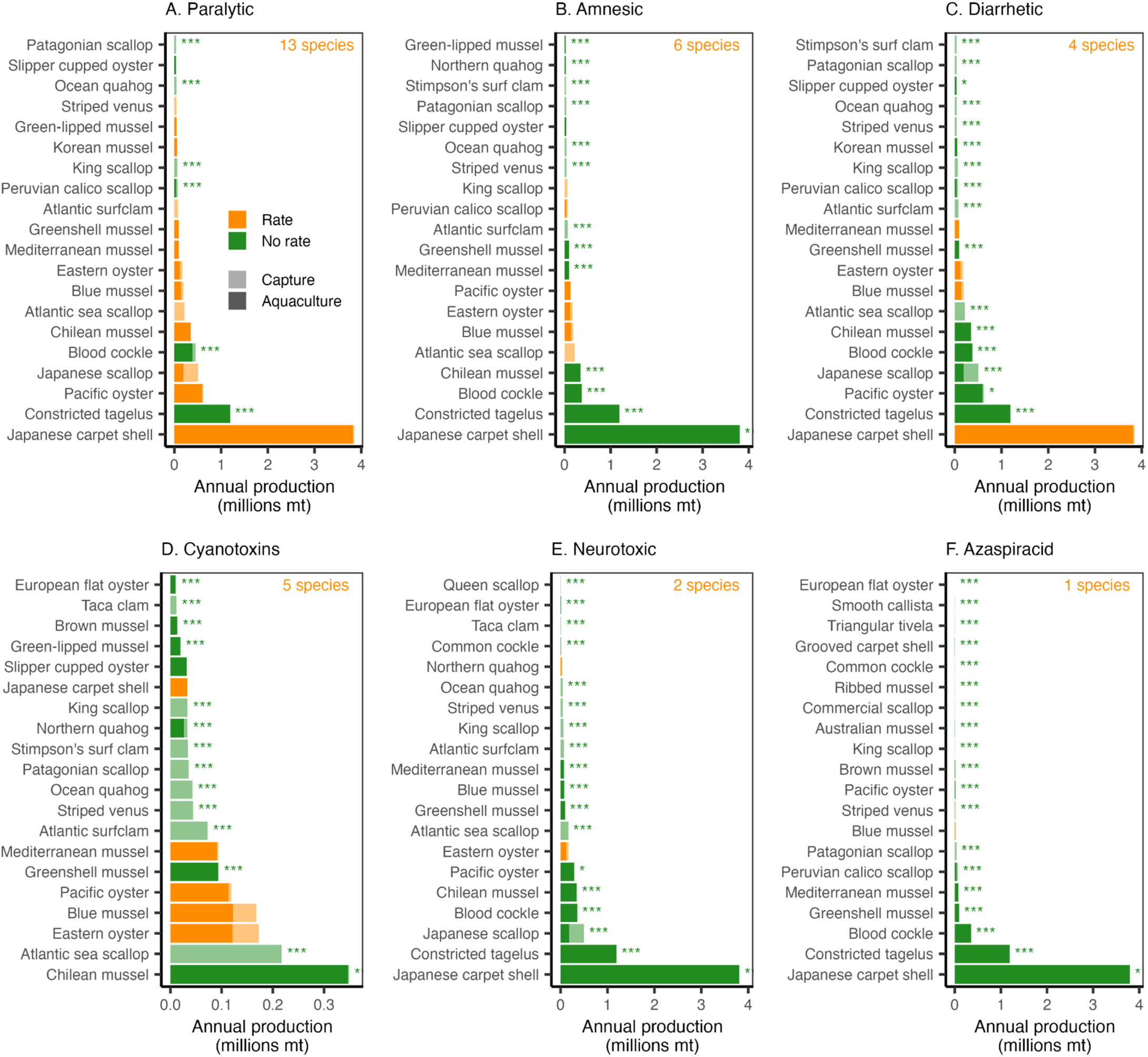
The twenty filter-feeding marine mollusc species with the greatest annual production (millions mt, 2014-2023) within Exclusive Economic Zones (**Fig. S5**) exposed to harmful algal blooms causing each of the evaluated biotoxin syndromes and an indication of whether (orange) or not (green) the depuration rate of the causative biotoxin has been studied. The species are sorted in order of decreasing annual production across both aquaculture (solid) and fisheries (transparent). The number of species whose depuration rates have been studied is printed in the top-right corner of each plot. The asterisks mark species whose depuration rate would contribute a new order (***), family (**), or genus (*) if studied. Ciguatera is evaluated separately because it is more strongly associated with large, predatory, tropical reef fish (**Fig. S7**).

Only six of the top-20 most harvested marine filter-feeding mollusc species from countries exposed to domoic acid have had their domoic acid depuration rates studied (**Fig. 5B**). Notably, none of the four most intensely harvested species have had their domoic acid depuration rates examined: Manila clam, constricted tagelus, blood cockle (*Tegillarca granosa*), or Chilean mussel (*Mytilus chilensis*). Domoic acid depuration rates for 1 new order, 4 new families, and 2 new genera could be gained through the evaluation of 11 of the 14 unstudied species (**Table S4**). In particular, evaluation of Manila clam (*Ruditapes philippinarum*), striped venus (*Chamelea gallina*), or northern quahog (*Mercenaria mercenaria*) would provide information for 22 harvested and 428 total species in the Veneridae family.

Only four of the top-20 most harvested marine filter-feeding mollusc species from countries exposed to DSTs have had their DST depuration rates studied (**Fig. 5C**). Notably, six of the seven most intensely harvested species have had their DST depuration rates examined: constricted tagelus, Pacific oyster (*Magallana gigas*), Japanese scallop (*Mizuhopecten yessoensis*), blood cockle, Chilean mussel, and Atlantic sea scallop (*Placopecten magellanicus*). DST depuration rates for 2 new orders, 3 new families, and 7 new genera could be gained through the evaluation of 14 of the 17 unstudied species (**Table S4**). In particular, evaluation of blood cockle would provide information for 8 harvested and 360 total species in the Arcoida order.

Only five of the top-20 most harvested marine filter-feeding mollusc species from countries exposed to cyanotoxins have had their cyanotoxin depuration rates studied (**Fig. 5D**). Notably, neither of the two most intensely harvested species have had their cyanotoxin depuration rates examined: Chilean mussel or Atlantic sea scallop. Cyanotoxin depuration rates for 4 new families, and 2 new genera could be gained through the evaluation of 14 of the 16 unstudied species (**Table S4**). In particular, evaluation of Atlantic sea scallop (*Placopecten magellanicus*), Patagonian scallop (*Zygochlamys patagonica*), or king scallop (*Pecten maximus*) would provide information for 20 harvested and 297 total species in the Pectinidae family.

Only two of the top-20 most harvested marine filter-feeding mollusc species from countries exposed to brevetoxins have had their brevetoxin depuration rates studied (**Fig. 5E**). Notably, none of the six most intensely harvested species have had their brevetoxin depuration rates examined: Manila clam, constricted tagelus, Japanese scallop, blood cockle, Chilean mussel, or Pacific oyster. Brevetoxin depuration rates for 2 new orders, 5 new families, and 5 new genera could be gained through the evaluation of 17 of the 18 unstudied species (**Table S4**). In particular, evaluation of Japanese scallop (*Mizuhopecten yessoensis*), Atlantic sea scallop, king scallop, or queen scallop (*Aequipecten opercularis*) would provide information for 20 harvested and 297 total species in the Pectinidae family.

Only one of the top-20 most harvested marine filter-feeding mollusc species from countries exposed to azaspiracids have had their azaspiracid depuration rates studied (**Fig. 5F**). Notably, none of the seven most intensely harvested species have had their azaspiracid depuration rates examined: Manila clam, constricted tagelus, blood cockle, greenshell mussel (*Perna canaliculus*), Mediterranean mussel (*Mytilus galloprovincialis*), Peruvian calico scallop (*Argopecten purpuratus*), or Patagonian scallop (*Zygochlamys patagonica*). Azaspiracid depuration rates for 4 new orders and 2 new genera could be gained through the evaluation of 17 of the 19 unstudied species (**Table S4**). In particular, evaluation of Manila clam, constricted tagelus, striped venus, common cockle (*Cerastoderma edule*), grooved carpet shell (*Ruditapes decussatus*), triangular tivela (*Tivela mactroides*), or smooth callista (*Callista chione*) would provide information for 58 harvested and 2549 total species in the Veneroida order.

Striped mullet (*Mugil cephalus*) is the only finfish in the top-50 most harvested finfish with known or hypothesized vulnerability to ciguatera from countries exposed to ciguatera to have had its ciguatera depuration rates studied (Ledreux et al., 2014) (**Fig. S7**). Otherwise, ciguatera depuration rates have only been quantified for minor fisheries species. Pinfish (*Lagodon rhomboides*) (Bennett and Robertson, 2021), orange-spotted grouper (*Epinephelus coioides*) (Li et al., 2020), and lionfish (*Pterois volitans*) (Leite et al., 2021) have had depuration rates measured but are the 88^th^, 99^th^, and 114^th^ most important large, predatory, reef-associated fisheries species occurring in countries where ciguatera is known to occur.

Although only 34 non-bivalves species have had their depuration rates studied, many non-bivalves have been observed within toxicities exceeding action thresholds (**Fig. S8**), highlighting the importance of studying biotoxin depuration in more non-bivalve species. In particular, a large number of harvested gastropods and crustaceans have been observed with PST toxicities far above the PST action threshold.

### 3.3 Predicting bivalve PST depuration rates

All four models exhibited good convergence (R^ < 1.01, no divergent transitions) and passed posterior predictive checks (**Figure S9**). The models with taxonomic random effects significantly outperformed the model with only life history, location, and tissue fixed effects (ΔELPD > 26; ΔR2=0.18; **Table S5**). The models with taxonomic random effects exhibited similar performance with the greatest statistical support for the taxonomically random effects model (ELPD=-229.4, R^2^=0.67) followed by the non-phylogenetic random effects model (ΔELPD=0.3, ΔELPD-SE=0.3, ΔR^2^=0.01) then the phylogenetic random effects model (ΔELPD=1.3, ΔELPD-SE=0.0, ΔR^2^=0.02). The taxonomic random effects model was therefore selected as the best performing model.

The taxonomic random effects model estimated that depuration rates are significantly slower (smaller) in the field relative to the lab (**Fig. 6A; Fig. S10A**). On average, field depuration rates are 2.5 times slower than lab depuration rates. The model estimated non-significant impacts of the three life history variables, which exhibited a mixture of non-significant positive (growth rate) and negative (maximum length, preferred temperature) effects. The model estimated significant differences in PST depuration rates among bivalve tissues with the hepatopancreas and soft tissues exhibiting faster depuration rates than other tissues. The conditional effects of the fixed effects variable, which estimate the depuration rate with all other effects estimated at their average, are shown in **Fig. S10**. The greater statistical support for this model over the “fixed effects only” model indicates a significant impact of taxonomy on depuration rates with the relative impacts of taxonomy on depuration rates illustrated in **Fig. 6B**. The greater support of this model over the phylogenetic model suggests that, based on the current data, while depuration rates vary by clade, they do not vary based on the relatedness of clades (**Table S5**).

**Figure 6.**
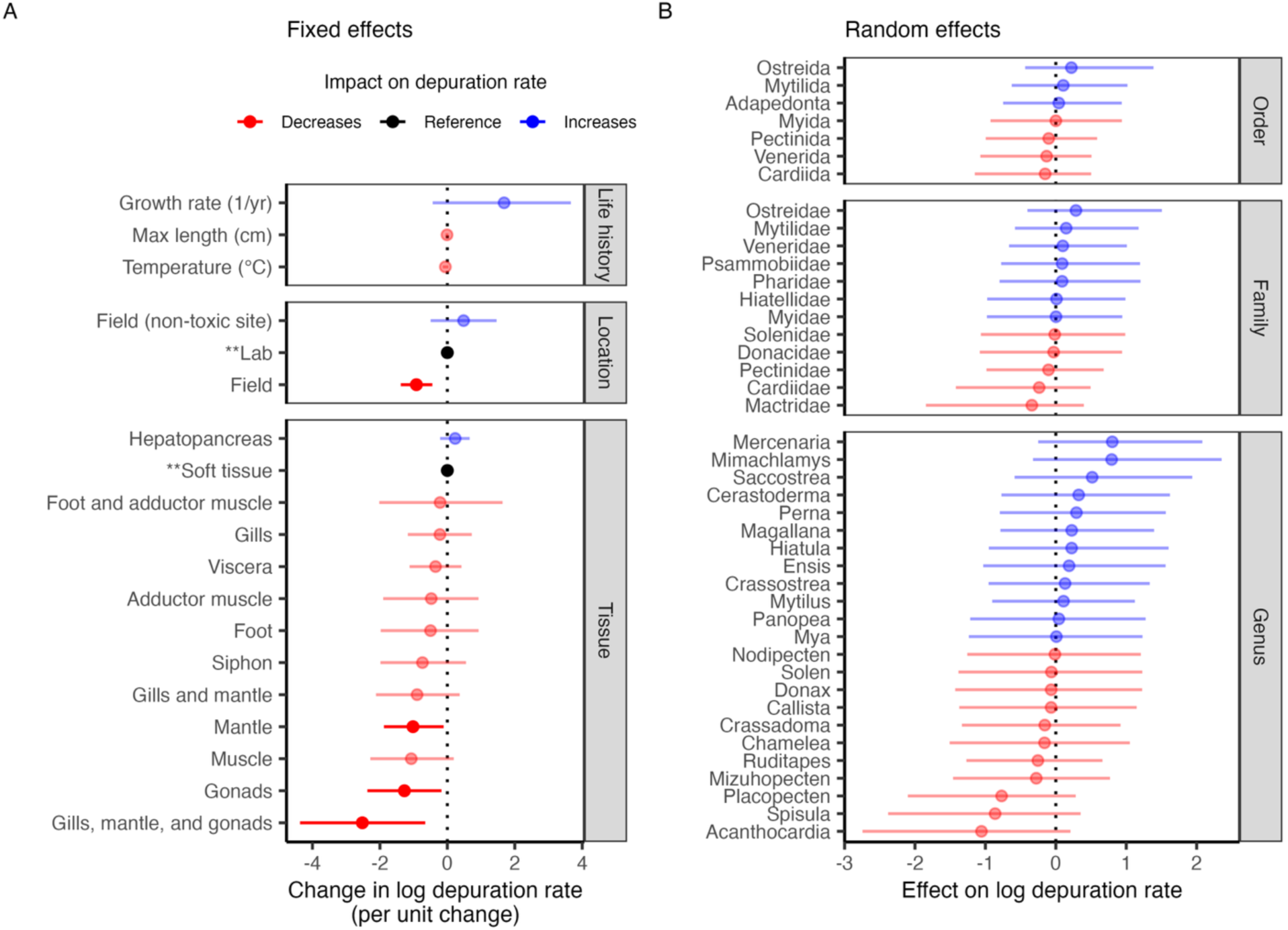
The **(A)** fixed effects and **(B)** random effects coefficients estimated by the final model. Points indicate the median estimate and lines indicate the 95% credible interval. Color indicates the impact of the point estimate of the effect on depuration rates. Transparency indicates whether the 95% credible intervals overlap with zero (solid=no overlap, transparent=overlap). The categorical fixed effects (location and tissue) are estimated relative to a reference level, which is marked with asterisks and is indicated in black. The conditional effects of the fixed effects variables are shown in **Fig. S10**.

The field-based PST depuration rates for species included in model fitting were estimated to vary from 0.0009 day^-1^ for smooth clam (*Callista chione*) to 0.20 day^-1^ for noble scallop (*Mimachlamys crassicostata*) (**Fig. 7; Table S6**). The field-based PST depuration rates for all harvested bivalves were estimated to vary from 0.0009 day^-1^ for smooth clam to 1.94 day^-1^ for Pacific calico scallop (*Argopecten ventricosus*) (**Fig. 8; Table S7**), after excluding exceptionally high and uncertain estimated depuration rates for giant clam (*Tridacna gigas*) and mangrove cupped oyster (*Crassostrea rhizophorae*). Giant clam (91.5 day^-1^) and mangrove copper oyster (16.7 day^-1^) exhibited high depuration rates because of their exceptionally large maximum size (137 cm) and fast growth rate (2.8 yr^-1^), respectively. Because no species in the Arcida or Liminda order were available in the training dataset, predictions for species in this order are not informed by taxonomic information. The lab- and field-based depuration rates predicted for harvested marine bivalves are provided in **Table S7**.

**Figure 7.**
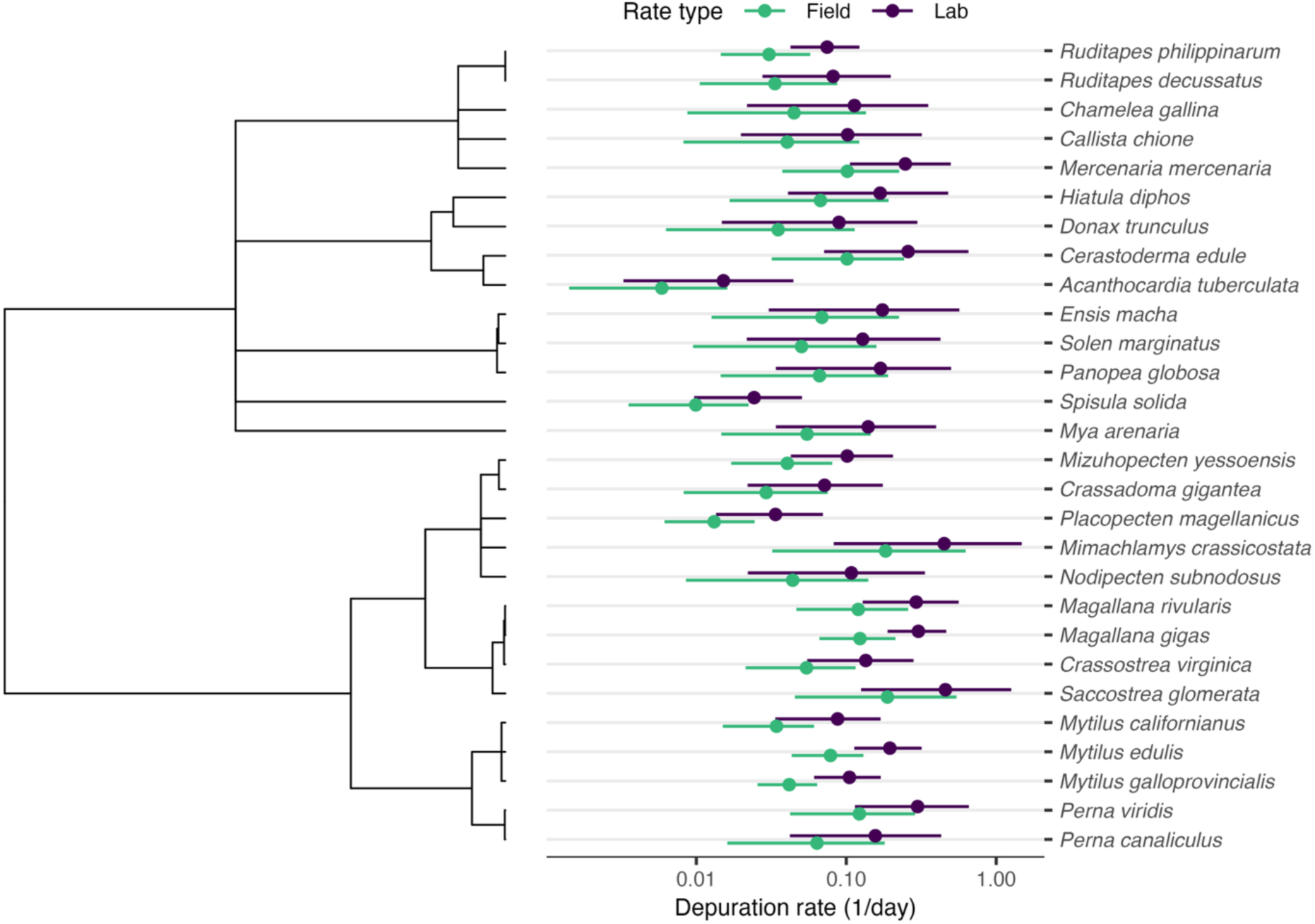
Paralytic shellfish toxin (PST) depuration rate estimates for species with measured depuration rates from the final regression model. Species are organized by phylogeny. Points indicate the median estimate and lines indicate the 95% credible interval. See **Table S6** for details.

**Figure 8.**
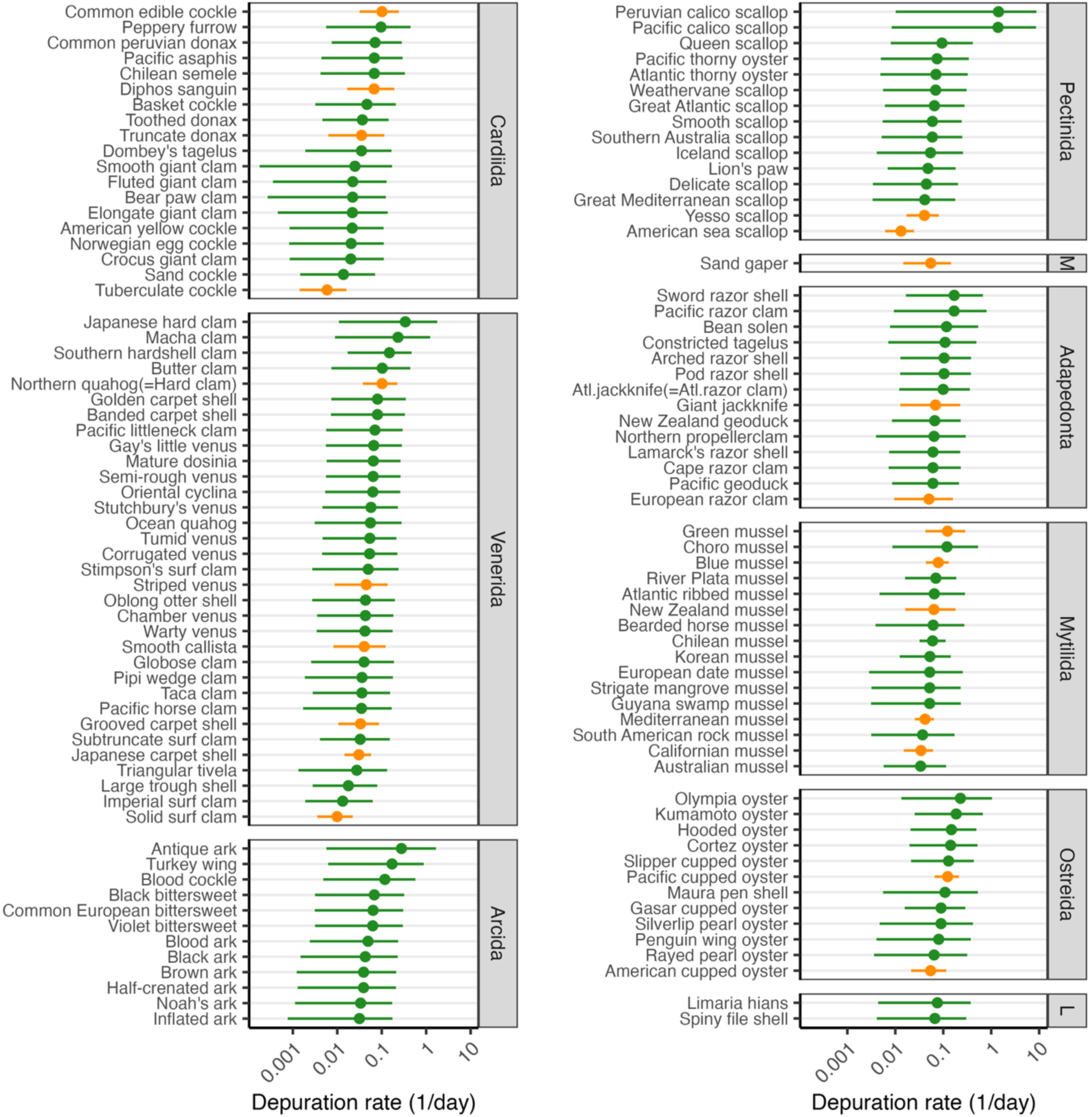
Paralytic shellfish toxin (PST) depuration rates for harvest marine bivalve species predicted by the final regression model. Species are organized by taxonomic order (M=Myida, L=Limida). Points indicate the median estimate and lines indicate the 95% credible interval. Species represented in the training data are marked in blue. Large predicted depuration rates for giant clam (*Tridacna gigas*) and mangrove cupped oyster (*Crassostrea rhizophorae*) are suppressed to ease visualization. See **Table S7** for details on the predicted depuration rates.

## 4. Discussion

Ensuring the safety of marine seafood is a daunting task. An estimated ∼200 taxa produce biotoxins that accumulate in marine food webs and threaten public health (Lundholm et al., 2009). These taxa produce a diverse range of toxins, which require independent chemical assays to detect and monitor (Hallegraeff et al., 2004). With over 3,000 species of marine fish and invertebrates harvested by commercial fisheries and aquaculture along hundreds of thousands of kilometers of coastline (FAO, 2024), it is impossible for biotoxin monitoring to cover all species, toxins, and locations. Thus, managers need every tool in the toolbox to design cost-effective biotoxin monitoring programs that protect public health while also limiting closures to commercially, recreationally, and culturally important coastal food systems.

Forecasting depuration timelines using depuration rates offers one such tool. With knowledge of how quickly a species eliminates toxins, managers can predict when toxicity will fall to levels deemed safe for human consumption. This information is valuable for several reasons. First, it provides seafood businesses with predictions of how long production is likely to be delayed, which can empower adaptive business decisions. For example, during long delays, harvesters may decide to pivot to new species, fishing grounds, or even alternative livelihoods (Moore et al., 2020). Second, such predictions could help managers decide whether to close fisheries, which could be preferred during short depuration timelines, or let them operate under an “evisceration order” that requires the removal of toxic viscera before sale, which can lower product value (Hackett et al., 2003), but may be preferable to closures during long depuration timelines. Finally, as described in the introduction, such predictions could be used to improve the efficiency of biotoxin monitoring (Blanco, 2009). Eliminating unnecessary tests would free up limited resources to allow monitoring of more sites or species, which has large economic and public health benefits. Monitoring more sites increases the resolution of biotoxin management and could lead to less restrictive fisheries closures (Free et al., 2022) while monitoring previously unmonitored species could more fully protect public health (Costa et al., 2017).

Our results show that depuration is slower in the field than in the lab, which has important implications for the application of lab-based depuration rates in natural settings. While lab studies quantify depuration rates while organisms are either staved or fed a non-toxic diet, field studies quantify depuration while organisms continue the uptake of biotoxins from dwindling blooms (Rourke et al., 2021), residual biotoxins in the food web (Yang et al., 2016), or dormant benthic cysts, which can be directly consumed by detritivores or planktivores if resuspended (Persson et al., 2006). This means that depuration timelines projected using lab-base depuration rates are likely to be optimistic (i.e., forecast faster declines in toxicity than reality). Fortunately, this is a favorable bias as testing sooner than necessary protects public health without unnecessarily delaying openings to fisheries or aquaculture operations (Free et al., 2022). The opposite bias (slower declines than reality) would be less favorable as it could delay testing until after toxicity has fallen below the action threshold, resulting in unnecessarily long closures to fisheries and aquaculture operations. However, managers may want to assume slower depuration rates than predicted by lab studies as more accurate forecasts will result in higher cost savings while still protecting public health and avoiding unnecessary closures. For example, managers could assume that depuration occurs 2.5 times more slowly in the field, the average estimated by our best performing Bayesian regression model.

Managers may also adjust the expected depuration rate and associated cadence of monitoring based on observations of temperature, food availability, body condition, or other factors that impact depuration rates. For example, managers may expect faster than average depuration rates if temperatures are warm (Bricelj et al., 2014), food availability and consumption is high (Bricelj and Shumway, 1998), or body condition is poor (Castberg et al., 2004). However, caution is necessary as these factors can have confounding impacts on bloom toxicity and biotoxin uptake. If warm water stimulates HAB productivity and toxicity (McKibben et al., 2017), elevated uptake may offset accelerated depuration. Similarly, if consumption is high due to either high food availability or low body condition, then rapid toxin uptake may offset accelerated depuration. Adjusting depuration rates based on these considerations may therefore be most useful once there is evidence that the bloom has abated and there is significantly less biotoxin available for uptake. Then, managers can use knowledge of recent temperature, food availability, or body condition to adjust the expected depuration rate and recalculate the optimized cadence of biotoxin monitoring.

Our review indicates that aquaculture farmers can take several actions to accelerate biotoxin depuration. Because depuration rates increase with food ingestion and egestion (Bricelj and Shumway, 1998), farms can enhance depuration by increasing food availability and/or stimulating food consumption. In the case of fed aquaculture for finfish and crustaceans, this could involve providing more food than usual. In the case of unfed bivalve aquaculture, consumption could be stimulated by reducing stocking density to reduce competition for ambient food (Cubillo et al., 2012). Another strategy could be to limit or inhibit feeding during a bloom to avoid biotoxin accumulation but also to reduce body condition and increase hunger to stimulate rapid consumption and depuration after the bloom abates (Castberg et al., 2004). Because depuration rates increase with temperature, which increases metabolism, consumption, and egestion (Bricelj et al., 2014; Novaczek et al., 1992), farmers could move racks, lines, rafts, or nets into warmer areas after a HAB to accelerate depuration. Furthermore, if warm water does not result in increased mortality, disease, HAB risk, or other detrimental effects, aquaculture could be strategically sited in warm areas to enhance depuration when necessary. Relatedly, aquaculture species with fast depuration rates (and slow uptake rates) may be good candidates for areas with high HAB risk. Finally, there is evidence that a number of industrial additives can be used to enhance depuration rates (Bian et al., 2024; Martinez-Albores et al., 2020), but these are often cost prohibitive or not allowed by regulation (Leal and Cristiano, 2024).

Biotoxin depuration rates of marine seafood species are understudied, especially for non-filter feeders. Higher trophic level species, including herbivorous gastropods, detritivorous crustaceans, and carnivorous fish, are particularly understudied, despite observations of unsafe levels of toxicity in these species (Costa et al., 2017; Deeds et al., 2008; Lefebvre and Robertson, 2010). The quantification of depuration rates for vulnerable gastropods, crustaceans, and finfish and their broader inclusion in biotoxin monitoring programs, where they are often overlooked (Costa et al., 2017), is thus important for fully protecting public health. It is frequently asserted that biotoxins in finfish pose minimum human health risk because the toxins do not accumulate in muscle tissue (e.g., Deeds et al., 2008; Lefebvre et al., 2002); however, the consumption of non-muscle tissue is common in many cultures and many preparation techniques use the whole fish (Golden et al., 2021). Furthermore, the processing of planktivorous forage fish into fish meal and fish oil uses the whole fish, presenting a potential pathway for unmonitored toxins to enter aquaculture and livestock feed (Adeyemo-Eleyode et al., 2025). Finally, the study of species that are not harvested by humans but represent important nodes in marine food webs is critical to understanding the trophic transfer of biotoxins (Holmes and Lewis, 2022; Lefebvre et al., 2002). Besides three unharvested copepod species (*Calanus finmarchicus* are harvested in Norway; FAO, 2024), we found that only two unharvested species (opossum shrimp, *Neomysis awatschensis* and Pacific mole crab, *Emerita analoga*) have had their depuration rates studied.

Although bivalves have undergone the most study, depuration rates have still not been quantified for most bivalve species harvested in commercial fisheries and aquaculture. For example, PST depuration rates are unquantified for 104 species, 60 entire genera, 12 entire families, and 2 entire orders of harvested marine bivalves. To our knowledge, we provide the first evidence that PST depuration rates are structured by bivalve taxonomy, meaning that closely related species exhibit similar depuration rates. Although we did not find evidence that PST depuration rates are phylogenetically conserved, this may be due to low taxonomic representation in current depuration studies, given that nested taxonomic identity was found to be an important predictor of PST depuration rates. As a result, managers can use either our predicted depuration rates or collated observed depuration rates from related species as proxies for species without known depuration rates. This finding also highlights the immense value of empirically quantifying PST depuration rates for a new order, family, or genus to provide guidance on depuration rates for the greatest number of new species. The expanded quantification of depuration rates would be useful to confirm whether depuration rates are taxonomically structured, or even phylogenetically conserved, across other toxins and taxonomic classes (i.e., not just a result for PSTs in bivalves). We suspect this is the case given the similarity in physiology and feeding ecology among related organisms (Leahy et al., 2025).

Biotoxin monitoring programs present a direct and underutilized data source for measuring depuration rates under natural conditions. Although marine biotoxin monitoring programs are used by many wealthy coastal countries (Andersen et al., 2004), our literature review found only 18 papers that directly reported depuration rates quantified from monitoring programs. These programs document the rise and fall of biotoxins in diverse species across many sites and years, representing a large and informative store of already collected data on seafood toxicokinetics. The analysis of these data would yield depuration rates for new species and toxins (e.g., Mcguire et al. (2025) provide the only source of DST depuration rates for eastern oyster, *Crassostrea virginica*) and would quantify the variability in depuration rates stemming from different environmental conditions (e.g., Blanco et al. (1997) show that impacts of salinity, temperature, light availability, and primary productivity on PST depuration in mussels), all without additional field or lab costs. As a start, we quantified depuration rates for 45 species-toxin combinations from 30 field studies that did not directly estimate depuration rates but were discovered in our systematic review due to the inclusion of depuration-related words in their abstract or keywords (e.g., Haya et al., 2003; Kvrgić et al., 2022; Rourke et al., 2021). However, this still overlooks the likely large number of biotoxin monitoring programs that are either not documented, are documented without these keywords, or are only documented in the grey or non-English literature. The optimization of monitoring programs using depuration rates derived from their own data would free up resources to expand monitoring to new species, thereby funding the iterative derivation of depuration rates and optimization of monitoring.

The utility of new depuration studies would be maximized through the adoption of a few best practices. In particular, surprisingly few of the evaluated studies (41% of papers) directly quantified depuration rates using standard depuration models, which complicates comparisons between studies and limits utility to managers. New studies should use standard depuration models (see Blanco, 2009) to quantify depuration rates to ease interpretability and use. If comparing one- and multi-compartment models, we recommend the use of Akaike information criterion (AIC) to select the most parsimonious model, recognizing that multi-compartment models have more parameters and should nearly always generate tighter, though not necessarily more parsimonious, fits (Burnham and Anderson, 2004). All of the studies comparing one- and multi-compartment models compared fits used either *R^2^* (e.g., Nielsen et al., 2016), which is biased to favor more complex models, or subjective judgements of fit and parsimony (e.g., Kennedy et al., 1992), which is not replicable. Next, given that depuration in real-world fisheries and aquaculture settings is likely to occur while feeding, new studies should always include a scenario in which the treated organism is fed during the depuration phase.

24% of the evaluated studies only considered depuration under starved conditions (e.g., Duinker et al., 2007b), limiting their relevance to real-world settings. Finally, depuration studies should consider management-relevant tissues to maximize utility. For example, 10% of the evaluated studies did not generate a depuration rate for tissues targeted for consumption by people or with the greatest toxin burden (e.g., focused only on the hemolymph; Schultz et al., 2008).

As the public health risk posed by HABs worsens under the combined effects of eutrophication and climate change (IPCC, 2019), managers will need every tool in the toolbox to improve the cost effectiveness of biotoxin monitoring. The use of depuration rates to adjust the cadence of monitoring during the depuration phase of HAB events is only one such tool. For example, the development of reliable early warning indicators of HAB risk could be used to more efficiently time the onset of biotoxin monitoring, sparing the need for testing while risk is low (Anderson et al., 2019). Similarly, knowledge of biotoxin uptake rates by seafood species could be used to optimize the cadence of testing during the uptake phase of HAB events. These advances will be critical to fully protecting public health while simultaneously avoiding unnecessary fisheries and aquaculture closures in a rapidly changing ocean.

## Acknowledgements

This work is the result of research funded by NOAA’s National Centers for Coastal Ocean Science Competitive Research Program, Climate Program Office, Ocean Acidification Program, and the U.S. Integrated Ocean Observing System Office under award NA22NOS4780171 to Oregon State University, and to NOAA’s Pacific Marine Environmental Laboratory.

