## Supplemental Materials for "Marine biotoxin depuration rates: management applications, research priorities, and predictions for unstudied species"

### Supplemental Tables and Figures

#### Supplemental Tables

**Table S1.** Attributes of papers that conduct studies of biotoxin depuration for aquatic organisms collected for the literature review analysis.

| Column | Description | Details |
| --- | --- | --- |
| paper_id | Paper id | Example: Lund et al. (1997) |
| comm_name | Common name | Example: Dungeness crab |
| sci_name | Scientific name | Example: Metacarcinus magister |
| syndrome | Syndrome | Amnesic, Diarrhetic, Paralytic, Azaspiracid, Ciguatera, Neurotoxic, Cyanotoxin |
| hab_species | HAB species | Examples: Pseudo-nitzschia australis |
| biotoxin | Biotoxin | Domoic acid, DSTs,PSTs, Azaspiracid, Ciguatoxin, Brevetoxin, Cyanotoxin, Other |
| subtoxin | Sub-biotoxin | Examples: dcSTX, GTX3, okadaic acid, etc |
| study_type | Study type | Llab, field, field (non-toxic) |
| exp_type | Experiment type | Examples: Temperature, Exposure, Diet type, etc. |
| treatment | Experimental treatment | Examples: 18°C, 20°C, 30 psu, 35 psu, etc. |
| feed_scenario | Feeding scenario | Fed clean, starved, ambient seawater, wild |
| tissue | Tissue type | Examples: digestive gland, hepatopancreas |
| source | Source of depuration rate | Examples: Figure 1, Table 1, Supplemental Data 1, Results 3.2 |
| rate_type | Derivation of depuration rate | provided, derived, digitized |
| ncomp | Number of compartments | None, zero, one, two, one vs. two, one vs. two vs. three |
| rate_hr | Depuration rate (1/hr) | |
| hlife_hr | half-life (hr) | |
| rate_d | Depuration rate (1/day) | |
| hlife_d | half-life (day) | |
| notes | General notes | |

**Table S2.** Tissue harmonization by class.

| Harmonized tissue | Synonyms |
|---|---|
| *Malacostraca* | |
| Hepatopancreas | Hepatopancreas, digestive gland |
| Soft tissue | Soft tissue, whole |
| *Gastropoda* | |
| Foot | Foot, muscle |
| Hepatopancreas | Hepatopancreas, digestive gland |
| *Bivalvia* | |
| Hepatopancreas | Hepatopancreas, digestive gland |
| Soft tissue | Soft tissue, tissue, edible tissue, edible portion, meat, flesh, whole flesh, whole, whole tissue, total tissue, non-viscera |

**Table S3.** The number of papers providing depuration rates by species and syndrome (par=paralytic; amn=amnesic; dia=diahhretic; cya=cyanotoxin; neu=neurotoxic; cig=ciguatera; aza=azaspiracid; oth=other).

| Class | Family | Scientific name | Common name | Par | Amn | Dia | Cya | Neu | Cig | Aza | Oth |
|---|---|---|---|---|---|---|---|---|---|---|---|
| Ascidiacea | Pyuridae | *Pyura chilensis* | Red sea squirt | 1 | | | | | | | |
| Bivalvia | Cardiidae | *Acanthocardia tuberculata* | Rough cockle | 1 | | | | | | | |
| Bivalvia | Cardiidae | *Cerastoderma edule* | Common cockle | 2 | | | | | | | |
| Bivalvia | Cyrenidae | *Corbicula fluminea* | Asian clam | | | | 1 | | | | |
| Bivalvia | Donacidae | *Donax trunculus* | Abrupt wedge shell | 1 | | 1 | | | | | |
| Bivalvia | Hiatellidae | *Panopea globosa* | Geoduck clam | 1 | | | | | | | |
| Bivalvia | Mactridae | *Spisula solida* | Surf clam | 1 | | | | | | | |
| Bivalvia | Mactridae | *Spisula solidissima* | Atlantic surfclam | 1 | | | | | | | |
| Bivalvia | Mesodesmatidae | *Paphies australis* | Pipi clam | | | | | | | | 1 |
| Bivalvia | Myidae | *Mya arenaria* | Soft-shell clam | 1 | 1 | | | | | | |
| Bivalvia | Mytilidae | *Aulacomya atra* | Chilean ribbed mussel | 1 | | | | | | | |
| Bivalvia | Mytilidae | *Geukensia demissa* | Atlantic ribbed mussel | | | 1 | | | | | 1 |
| Bivalvia | Mytilidae | *Mytella guyanensis* | Trinidad swamp mussel | | | 1 | | | | | |
| Bivalvia | Mytilidae | *Mytilus californianus* | California mussel | 1 | 1 | | 1 | | | | |
| Bivalvia | Mytilidae | *Mytilus chilensis* | Chilean mussel | 3 | | | | | | | 1 |
| Bivalvia | Mytilidae | *Mytilus coruscus* | Korean hard-shelled mussel | 1 | | | | | | | |
| Bivalvia | Mytilidae | *Mytilus edulis* | Blue mussel | 7 | 5 | 10 | 6 | | | 1 | 6 |
| Bivalvia | Mytilidae | *Mytilus galloprovincialis* | Mediterranean mussel | 9 | | 5 | 3 | | | | |
| Bivalvia | Mytilidae | *Mytilus spp.* | Mussels | 1 | | | | | | 1 | 1 |
| Bivalvia | Mytilidae | *Perna canaliculus* | Green-lipped mussel | 1 | | | | | | | 1 |
| Bivalvia | Mytilidae | *Perna perna* | Brown mussel | | | 1 | | | | | |
| Bivalvia | Mytilidae | *Perna viridis* | Asian green mussel | 3 | | 1 | | 1 | | | |
| Bivalvia | Ostreidae | *Crassostrea tulipa* | West Africa mangrove oyster | | | 1 | | | | | |
| Bivalvia | Ostreidae | *Crassostrea virginica* | Eastern oyster | 2 | 1 | 1 | 1 | 3 | | | 1 |
| Bivalvia | Ostreidae | *Magallana gigas* | Pacific oyster | 6 | 1 | | 4 | | | | 2 |
| Bivalvia | Ostreidae | *Magallana rivularis* | Jinjiang oyster | 2 | | | | | | | |
| Bivalvia | Ostreidae | *Ostrea chilensis* | Flat oyster | 1 | | | | | | | |
| Bivalvia | Ostreidae | *Ostrea edulis* | European flat oyster | | | 2 | | | | | 2 |
| Bivalvia | Ostreidae | *Saccostrea glomerata* | Sydney rock oyster | 1 | | | | | | | |
| Bivalvia | Pectinidae | *Aequipecten opercularis* | Queen scallop | | 1 | | | | | | 1 |
| Bivalvia | Pectinidae | *Argopecten irradians* | Bay scallop | 4 | | 1 | | | | | |
| Bivalvia | Pectinidae | *Argopecten purpuratus* | Peruvian calico scallop | | 1 | | | | | | |
| Bivalvia | Pectinidae | *Crassadoma gigantea* | Purple-hinged rock scallop | 1 | | 1 | | | | | |
| Bivalvia | Pectinidae | *Mimachlamys crassicostata* | Noble scallop | 2 | | | | | | | |
| Bivalvia | Pectinidae | *Mimachlamys varia* | Variegated scallop | | 1 | | | | | | |
| Bivalvia | Pectinidae | *Mizuhopecten yessoensis* | Japanese scallop | 4 | | | | | | | |
| Bivalvia | Pectinidae | *Nodipecten subnodosus* | Pacific giant lions-paw scallop | 1 | | | | | | | |
| Bivalvia | Pectinidae | *Pecten maximus* | King scallop | | 6 | | | | | | |
| Bivalvia | Pectinidae | *Placopecten magellanicus* | Atlantic sea scallop | 2 | 2 | | | | | | |
| Bivalvia | Pectinidae | *Scaeochlamys farreri* | Farrer's scallop | 2 | | | | | | 1 | |
| Bivalvia | Pharidae | *Ensis macha* | Navaja clam | 1 | | | | | | | |
| Bivalvia | Pharidae | *Siliqua patula* | Pacific razor clam | | 2 | | | | | | |
| Bivalvia | Pharidae | *Sinonovacula constricta* | Constricted tagelus | | | | | | | | 1 |
| Bivalvia | Psammobiidae | *Hiatula diphos* | Purple clam | 1 | | | | | | | |
| Bivalvia | Solenidae | *Solen marginatus* | Grooved razor shell | 1 | | | | | | | |
| Bivalvia | Veneridae | *Anomalocardia flexuosa* | Anomalocardia clam | | | 1 | | | | | |
| Bivalvia | Veneridae | *Callista chione* | Smooth clam | 1 | | | | | | | |
| Bivalvia | Veneridae | *Chamelea gallina* | Warty venus | 1 | | | | | | | |
| Bivalvia | Veneridae | *Mercenaria campechiensis* | Hard clam | | | | | 1 | | | |
| Bivalvia | Veneridae | *Mercenaria mercenaria* | Northern quahog | 1 | | | | 1 | | | 2 |
| Bivalvia | Veneridae | *Ruditapes decussatus* | Grooved carpet shell | 1 | | 1 | | | | | 2 |
| Bivalvia | Veneridae | *Ruditapes philippinarum* | Manila clam | 3 | | 1 | 1 | | | | |
| Cephalopoda | Octopodidae | *Octopus vulgaris* | Common octopus | 1 | | | | | | | |
| Copepoda | Acartiidae | *Acartia clausi* | A. clausi copepod | 2 | 1 | | | | | | |
| Copepoda | Acartiidae | *Acartia hudsonica* | A. hudsonica copepod | 1 | | | | | | | |
| Copepoda | Calanidae | *Calanus finmarchicus* | C. finmarchicus copepod | | 2 | | | | | | |
| Copepoda | Calanidae | *Calanus glacialis* | C. glacialis copepod | | 1 | | | | | | |
| Copepoda | Temoridae | *Eurytemora affinis* | E. affinis copepod | | | | 1 | | | | |
| Dinophyceae | Noctilucaceae | *Noctiluca scintillans* | N. scintillans dinoflagellate | 1 | | | | | | | |
| Gastropoda | Haliotidae | *Haliotis midae* | South African abalone | 1 | | | | | | | |
| Gastropoda | Haliotidae | *Haliotis rubra* | Australian blacklip abalone | 1 | | | | | | | |
| Gastropoda | Viviparidae | *Sinotaia quadrata* | Chinese freshwater snail | | | | 1 | | | | |

| | | | | | | | | | | |
|---|---|---|---|---|---|---|---|---|---|---|
| Malacostraca | Callichiridae | *Callichirus major* | Carolinian ghost shrimp | | | 1 | | | | |
| Malacostraca | Cancridae | *Cancer pagurus* | Brown crab | | | 2 | | | | 2 |
| Malacostraca | Cancridae | *Metacarcinus magister* | Dungeness crab | | 3 | | | | | |
| Malacostraca | Hippidae | *Emerita analoga* | Pacific mole crab | 1 | | | | | | |
| Malacostraca | Mysidae | *Neomysis awatschensis* | Mysid crustacean | | | | 1 | | | |
| Malacostraca | Palinuridae | *Jasus edwardsii* | Southern rock lobster | 3 | | | | | | |
| Malacostraca | Palinuridae | *Panulirus stimpsoni* | Chinese spiny lobster | 1 | | | | | | |
| Malacostraca | Penaeidae | *Penaeus monodon* | Black tiger prawn | | | | 1 | | | |
| Mammalia | Delphinidae | *Tursiops truncatus* | Bottlenose dolphin | | | | | 1 | | |
| Teleostei | Alosidae | *Brevoortia tyrannus* | Atlantic menhaden | | | | | 1 | | |
| Teleostei | Batrachoididae | *Opsanus beta* | Gulf toadfish | | | | | 1 | | |
| Teleostei | Ephippidae | *Chaetodipterus faber* | Atlantic spadefish | | | 1 | | | | |
| Teleostei | Epinephelidae | *Epinephelus coioides* | Orange-spotted grouper | | | | | | 1 | |
| Teleostei | Mugilidae | *Mugil cephalus* | Striped mullet | | | | | 2 | 1 | |
| Teleostei | Mugilidae | *Mugil liza* | Lebranche mullet | | | 1 | | | | |
| Teleostei | Salmonidae | *Oncorhynchus kisutch* | Coho salmon | | 1 | | | | | |
| Teleostei | Scorpaenidae | *Pterois volitans* | Lionfish | | | | | | 1 | |
| Teleostei | Sparidae | *Acanthopagrus schlegelii* | Black sea bream | 1 | | | | | | |
| Teleostei | Sparidae | *Diplodus sargus* | White seabream | 1 | | | | | | |
| Teleostei | Sparidae | *Lagodon rhomboides* | Pinfish | | | | | | 1 | |
| Teleostei | Sparidae | *Sparus aurata* | Gilthead seabream | 1 | | | | | | |
| Teleostei | Tetraodontidae | *Takifugu obscurus* | Obscure pufferfish | | | | | | | 1 |
| Thecostraca | Balanidae | *Austromegabalanus psittacus* | Giant barnacle | 1 | | | | | | |

**Table S4.** Diversity of orders, families, and genera of harvested marine molluscs.

| | | | | Production | Number of species | |
|---|---|---|---|---|---|---|
| Syndrome | Type | Group | Species | (1000s mt) | Harvested | Total |
| Amnesic | Order | Arcoida | Blood cockle (Tegillarca granosa) | 375.1 | 8 | 360 |
| | Family | Veneridae | Manila clam (Ruditapes philippinarum), Striped venus (Chamelea gallina), Northern quahog (Mercenaria mercenaria) | 3885.3 | 22 | 428 |
| | | Solecurtidae | Constricted tagelus (Sinonovacula constricta) | 1191.3 | 2 | 26 |
| | | Mactridae | Atlantic surfclam (Spisula solidissima), Stimpson's surf clam (Mactromeris polynyma) | 106.1 | 10 | 116 |
| | | Arcticidae | Ocean quahog (Arctica islandica) | 42.8 | 1 | 1 |
| | Genus | Perna | Greenshell mussel (Perna canaliculus), Green-lipped mussel (Perna viridis) | 118.9 | 3 | 5 |
| | | Zygochlamys | Patagonian scallop (Zygochlamys patagonica) | 35.5 | 2 | 1 |
| Azaspiracid | Order | Veneroida | Manila clam (Ruditapes philippinarum), Constricted tagelus (Sinonovacula constricta), Striped venus (Chamelea gallina), Common cockle (Cerastoderma edule), Grooved carpet shell (Ruditapes decussatus), Triangular tivela (Tivela mactroides), Smooth callista (Callista chione) | 5015.8 | 58 | 2549 |
| | | Arcoida | Blood cockle (Tegillarca granosa) | 356.8 | 8 | 360 |
| | | Ostreoida | Peruvian calico scallop (Argopecten purpuratus), Patagonian scallop (Zygochlamys patagonica), King scallop (Pecten maximus), Commercial scallop (Pecten fumatus), European flat oyster (Ostrea edulis) | 107.5 | 26 | 548 |
| | | Ostreida | Pacific oyster (Magallana gigas) | 16.3 | 5 | 88 |
| | Genus | Perna | Greenshell mussel (Perna canaliculus), Brown mussel (Perna perna) | 107.2 | 3 | 5 |
| | | Aulacomya | Ribbed mussel (Aulacomya atra) | 2.5 | 1 | 3 |
| Cyanotoxin | Family | Pectinidae | Atlantic sea scallop (Placopecten magellanicus), Patagonian scallop (Zygochlamys patagonica), King scallop (Pecten maximus) | 286.3 | 20 | 297 |
| | | Veneridae | Striped venus (Chamelea gallina), Northern quahog (Mercenaria mercenaria), Manila clam (Ruditapes philippinarum), Taca clam (Leukoma thaca) | 122.0 | 22 | 428 |
| | | Mactridae | Atlantic surfclam (Spisula solidissima), Stimpson's surf clam (Mactromeris polynyma) | 106.1 | 10 | 116 |
| | | Arcticidae | Ocean quahog (Arctica islandica) | 42.8 | 1 | 1 |
| | Genus | Perna | Greenshell mussel (Perna canaliculus), Green-lipped mussel (Perna viridis), Brown mussel (Perna perna) | 126.5 | 3 | 5 |
| | | Ostrea | European flat oyster (Ostrea edulis) | 10.6 | 3 | 20 |
| Diarrhetic | Order | Ostreida | Pacific oyster (Magallana gigas), Slipper cupped oyster (Magallana bilineata) | 657.5 | 5 | 88 |
| | | Arcoida | Blood cockle (Tegillarca granosa) | 375.7 | 8 | 360 |
| | Family | Solecurtidae | Constricted tagelus (Sinonovacula constricta) | 1191.3 | 2 | 26 |
| | | Mactridae | Atlantic surfclam (Spisula solidissima), Stimpson's surf clam (Mactromeris polynyma) | 106.1 | 10 | 116 |
| | | Arcticidae | Ocean quahog (Arctica islandica) | 42.8 | 1 | 1 |
| | Genus | Ruditapes | Manila clam (Ruditapes philippinarum) | 3848.4 | 2 | 4 |
| | | Mizuhopecten | Japanese scallop (Mizuhopecten yessoensis) | 505.8 | 1 | 1 |
| | | Placopecten | Atlantic sea scallop (Placopecten magellanicus) | 217.5 | 1 | 1 |
| | | Perna | Greenshell mussel (Perna canaliculus) | 93.5 | 3 | 5 |
| | | Pecten | King scallop (Pecten maximus) | 62.2 | 4 | 18 |
| | | Chamelea | Striped venus (Chamelea gallina) | 46.1 | 1 | 3 |
| | | Zygochlamys | Patagonian scallop (Zygochlamys patagonica) | 35.5 | 2 | 1 |
| Neurotoxic | Order | Arcoida | Blood cockle (Tegillarca granosa) | 357.1 | 8 | 360 |
| | | Ostreida | Pacific oyster (Magallana gigas) | 293.1 | 5 | 88 |
| | Family | Solecurtidae | Constricted tagelus (Sinonovacula constricta) | 1191.3 | 2 | 26 |
| | | Pectinidae | Japanese scallop (Mizuhopecten yessoensis), Atlantic sea scallop (Placopecten magellanicus), King scallop (Pecten maximus), Queen scallop (Aequipecten opercularis) | 734.1 | 20 | 297 |
| | | Mactridae | Atlantic surfclam (Spisula solidissima) | 71.8 | 10 | 116 |
| | | Arcticidae | Ocean quahog (Arctica islandica) | 42.8 | 1 | 1 |
| | | Cardiidae | Common cockle (Cerastoderma edule) | 14.3 | 13 | 222 |
| | Genus | Ruditapes | Manila clam (Ruditapes philippinarum) | 3813.6 | 2 | 4 |
| | | Mytilus | Chilean mussel (Mytilus chilensis), Blue mussel (Mytilus edulis), Mediterranean mussel (Mytilus galloprovincialis) | 503.5 | 7 | 12 |
| | | Chamelea | Striped venus (Chamelea gallina) | 45.7 | 1 | 3 |
| | | Leukoma | Taca clam (Leukoma thaca) | 11.9 | 2 | 3 |
| | | Ostrea | European flat oyster (Ostrea edulis) | 11.0 | 3 | 20 |
| Paralytic | Order | Arcoida | Blood cockle (Tegillarca granosa) | 453.6 | 8 | 360 |
| | Family | Solecurtidae | Constricted tagelus (Sinonovacula constricta) | 1191.3 | 2 | 26 |
| | | Arcticidae | Ocean quahog (Arctica islandica) | 42.8 | 1 | 1 |
| | Genus | Argopecten | Peruvian calico scallop (Argopecten purpuratus) | 70.2 | 3 | 9 |
| | | Pecten | King scallop (Pecten maximus) | 62.2 | 4 | 18 |
| | | Chamelea | Striped venus (Chamelea gallina) | 46.1 | 1 | 3 |
| | | Zygochlamys | Patagonian scallop (Zygochlamys patagonica) | 35.5 | 2 | 1 |

**Table S5.** Performance statistics used in model selection.

| **Model** | **ΔELPD** | **ELPD (sd)** | **LOOIC (sd)** | **P (sd)** | **$R^2$ (CI)** |
|---|---|---|---|---|---|
| Taxonomically nested random effects | 0 | -229.4 (9.2) | 458.8 (18.3) | 27 (2.6) | 0.67 (0.6-0.72) |
| Species random effects | -0.3 | -229.7 (9) | 459.4 (18.1) | 26.9 (2.5) | 0.66 (0.59-0.71) |
| Phylogenetic random effects | -1.3 | -230.8 (9.2) | 461.5 (18.5) | 25.1 (2.6) | 0.65 (0.57-0.71) |
| Fixed effects only | -26 | -255.4 (9.5) | 510.8 (19) | 16.5 (2.2) | 0.49 (0.4-0.56) |

* ELPD = expected log predictive density (higher = better); LOOIC = leave-one-out information criterion (lower = better); $R^2$ = Bayesian $R^2$ (higher = better); P = effective number of parameters; SE = standard error; CI = 95% confidence interval

**Table S6.** Field- and lab-based paralytic shellfish toxin depuration rates predicted by the best performing regression model for species included in model training along with the required predictor variables.

| Order | Family | Genus | Scientific name | Common name | Length (cm) | K (1/yr) | Temp (°C) | Field (95% CI) | Lab (95% CI) |
|---|---|---|---|---|---|---|---|---|---|
| Adapedonta | Pharidae | Ensis | *Ensis macha* | Navaja clam | 16.5 | 0.225 | 11 | 0.069 (0.013-0.225) | 0.174 (0.03-0.568) |
| Adapedonta | Hiatellidae | Panopea | *Panopea globosa* | Geoduck clam | 19.8 | 0.33 | 8 | 0.066 (0.015-0.19) | 0.169 (0.034-0.501) |
| Adapedonta | Solenidae | Solen | *Solen marginatus* | Grooved razor shell | 17 | 0.28 | 10 | 0.05 (0.009-0.159) | 0.129 (0.022-0.425) |
| Cardiida | Cardiidae | Acanthocardia | *Acanthocardia tuberculata* | Rough cockle | 9 | 0.669 | 26.5 | 0.006 (0.001-0.016) | 0.015 (0.003-0.044) |
| Cardiida | Cardiidae | Cerastoderma | *Cerastoderma edule* | Common cockle | 5.6 | 0.62 | 10 | 0.101 (0.032-0.243) | 0.258 (0.071-0.653) |
| Cardiida | Donacidae | Donax | *Donax trunculus* | Abrupt wedge shell | 4.4 | 0.7 | 26.5 | 0.035 (0.006-0.114) | 0.089 (0.015-0.298) |
| Cardiida | Psammobiidae | Hiatula | *Hiatula diphos* | Purple clam | 6 | 0.825 | 26.5 | 0.067 (0.017-0.192) | 0.168 (0.041-0.478) |
| Myida | Myidae | Mya | *Mya arenaria* | Soft-shell clam | 10 | 0.3 | 10 | 0.055 (0.015-0.146) | 0.14 (0.034-0.397) |
| Mytilida | Mytilidae | Mytilus | *Mytilus californianus* | California mussel | 25.5 | 0.25 | 15 | 0.034 (0.015-0.061) | 0.088 (0.034-0.169) |
| Mytilida | Mytilidae | Mytilus | *Mytilus edulis* | Blue mussel | 11 | 0.24 | 8 | 0.078 (0.043-0.13) | 0.195 (0.113-0.318) |
| Mytilida | Mytilidae | Mytilus | *Mytilus galloprovincialis* | Mediterranean mussel | 16.5 | 0.26 | 17 | 0.042 (0.026-0.064) | 0.105 (0.061-0.17) |
| Mytilida | Mytilidae | Perna | *Perna canaliculus* | Green-lipped mussel | 15 | 0.6 | 26 | 0.064 (0.016-0.18) | 0.156 (0.042-0.43) |
| Mytilida | Mytilidae | Perna | *Perna viridis* | Asian green mussel | 16.5 | 1.07 | 26 | 0.122 (0.042-0.287) | 0.299 (0.114-0.657) |
| Ostreida | Ostreidae | Crassostrea | *Crassostrea virginica* | Eastern oyster | 30 | 0.565 | 24 | 0.054 (0.021-0.115) | 0.135 (0.055-0.281) |
| Ostreida | Ostreidae | Magallana | *Magallana gigas* | Pacific oyster | 45 | 0.68 | 13 | 0.123 (0.066-0.213) | 0.303 (0.189-0.464) |
| Ostreida | Ostreidae | Magallana | *Magallana rivularis* | Jinjiang oyster | 20.88 | 0.84 | 20.5 | 0.12 (0.046-0.259) | 0.293 (0.129-0.562) |
| Ostreida | Ostreidae | Saccostrea | *Saccostrea glomerata* | Sydney rock oyster | 5 | 0.7625 | 24 | 0.188 (0.045-0.544) | 0.458 (0.125-1.26) |
| Pectinida | Pectinidae | Crassadoma | *Crassadoma gigantea* | Purple-hinged rock scallop | 25 | 0.5035 | 16 | 0.029 (0.008-0.075) | 0.072 (0.022-0.175) |
| Pectinida | Pectinidae | Mimachlamys | *Mimachlamys crassicostata* | Noble scallop | 9.49 | 0.6626 | 17 | 0.183 (0.032-0.626) | 0.45 (0.082-1.479) |
| Pectinida | Pectinidae | Mizuhopecten | *Mizuhopecten yessoensis* | Japanese scallop | 25 | 0.925 | 18 | 0.04 (0.017-0.081) | 0.101 (0.043-0.205) |
| Pectinida | Pectinidae | Nodipecten | *Nodipecten subnodosus* | Pacific giant lions-paw scallop | 15 | 0.5035 | 16 | 0.044 (0.009-0.14) | 0.108 (0.022-0.335) |
| Pectinida | Pectinidae | Placopecten | *Placopecten magellanicus* | Atlantic sea scallop | 20 | 0.28 | 8 | 0.013 (0.006-0.024) | 0.034 (0.014-0.07) |
| Venerida | Veneridae | Callista | *Callista chione* | Smooth clam | 11 | 0.24 | 12 | 0.04 (0.008-0.122) | 0.102 (0.02-0.319) |
| Venerida | Veneridae | Chamelea | *Chamelea gallina* | Warty venus | 5 | 0.415 | 14 | 0.045 (0.009-0.135) | 0.113 (0.022-0.352) |
| Venerida | Veneridae | Mercenaria | *Mercenaria mercenaria* | Northern quahog | 13 | 0.3257 | 17 | 0.102 (0.037-0.225) | 0.248 (0.106-0.498) |
| Venerida | Veneridae | Ruditapes | *Ruditapes decussatus* | Grooved carpet shell | 8 | 0.48 | 17 | 0.033 (0.011-0.087) | 0.082 (0.028-0.198) |
| Venerida | Veneridae | Ruditapes | *Ruditapes philippinarum* | Manila clam | 8 | 0.52 | 18 | 0.031 (0.015-0.058) | 0.075 (0.042-0.122) |
| Venerida | Mactridae | Spisula | *Spisula solida* | Surf clam | 5 | 0.3307 | 11 | 0.01 (0.004-0.022) | 0.024 (0.01-0.051) |

**Table S7.** Field- and lab-based paralytic shellfish toxin depuration rates predicted by the best performing regression model for all harvested bivalve species along with the required predictor variables.

| Order | Family | Genus | Scientific name | Common name | Length (cm) | K (1/yr) | Temp (°C) | Field (95% CI) | Lab (95% CI) |
|---|---|---|---|---|---|---|---|---|---|
| Adapedonta | Hiatellidae | Cyrtodaria | *Cyrtodaria siliqua* | Northern propellerclam | 11 | 0.07 | 8 | 0.064 (0.004-0.294) | 0.161 (0.01-0.75) |
| Adapedonta | Pharidae | Ensis | *Ensis ensis* | Pod razor shell | 16.5 | 0.4625 | 11 | 0.104 (0.013-0.376) | 0.261 (0.031-0.944) |
| Adapedonta | Pharidae | Ensis | *Ensis leei* | Atl.jackknife(=Atl.razor clam) | 20 | 0.4625 | 11 | 0.099 (0.012-0.357) | 0.25 (0.028-0.897) |
| Adapedonta | Pharidae | Ensis | *Ensis macha* | Giant jackknife | 16.5 | 0.225 | 11 | 0.069 (0.013-0.225) | 0.174 (0.03-0.568) |
| Adapedonta | Pharidae | Ensis | *Ensis magnus* | Arched razor shell | 16.5 | 0.4625 | 11 | 0.104 (0.013-0.379) | 0.261 (0.03-0.951) |
| Adapedonta | Pharidae | Ensis | *Ensis siliqua* | Sword razor shell | 15.5 | 0.7 | 11 | 0.169 (0.017-0.671) | 0.421 (0.039-1.644) |
| Adapedonta | Hiatellidae | Panopea | *Panopea generosa* | Pacific geoduck | 17.5 | 0.199 | 8 | 0.061 (0.009-0.213) | 0.154 (0.02-0.537) |
| Adapedonta | Hiatellidae | Panopea | *Panopea zelandica* | New Zealand geoduck | 17.5 | 0.25 | 8 | 0.066 (0.009-0.23) | 0.168 (0.02-0.594) |
| Adapedonta | Pharidae | Pharus | *Pharus legumen* | Bean solen | 9.982 | 0.4075 | 9 | 0.117 (0.008-0.533) | 0.291 (0.019-1.295) |
| Adapedonta | Pharidae | Siliqua | *Siliqua patula* | Pacific razor clam | 18 | 0.59 | 7 | 0.169 (0.009-0.798) | 0.422 (0.023-1.969) |
| Adapedonta | Pharidae | Sinonovacula | *Sinonovacula constricta* | Constricted tagelus | 15.5 | 0.4075 | 9 | 0.109 (0.007-0.49) | 0.272 (0.018-1.215) |
| Adapedonta | Solenidae | Solen | *Solen capensis* | Cape razor clam | 11 | 0.28 | 10 | 0.061 (0.007-0.231) | 0.153 (0.017-0.587) |
| Adapedonta | Solenidae | Solen | *Solen lamarckii* | Lamarck's razor shell | 10 | 0.28 | 10 | 0.061 (0.007-0.225) | 0.154 (0.018-0.574) |
| Adapedonta | Solenidae | Solen | *Solen marginatus* | European razor clam | 17 | 0.28 | 10 | 0.05 (0.009-0.159) | 0.129 (0.022-0.425) |
| Arcida | Arcidae | Anadara | *Anadara antiquata* | Antique ark | 10.5 | 1.54 | 31.75 | 0.278 (0.006-1.666) | 0.676 (0.014-3.854) |
| Arcida | Arcidae | Anadara | *Anadara broughtonii* | Inflated ark | 8 | 0.2359 | 31.75 | 0.031 (0.001-0.175) | 0.078 (0.002-0.422) |
| Arcida | Arcidae | Anadara | *Anadara kagoshimensis* | Half-crenated ark | 6.2 | 0.428 | 31.75 | 0.039 (0.001-0.209) | 0.096 (0.003-0.506) |
| Arcida | Arcidae | Anadara | *Anadara similis* | Brown ark | 8 | 0.428 | 31.75 | 0.039 (0.001-0.211) | 0.096 (0.003-0.516) |
| Arcida | Arcidae | Anadara | *Anadara tuberculosa* | Black ark | 8 | 0.515 | 31.75 | 0.043 (0.001-0.229) | 0.105 (0.004-0.552) |
| Arcida | Arcidae | Arca | *Arca noae* | Noah's ark | 7 | 0.165 | 26.5 | 0.034 (0.001-0.172) | 0.082 (0.003-0.419) |
| Arcida | Arcidae | Arca | *Arca zebra* | Turkey wing | 10 | 1.2 | 26.5 | 0.171 (0.006-0.882) | 0.416 (0.016-2.085) |
| Arcida | Glycymerididae | Glycymeris | *Glycymeris glycymeris* | Common European bittersweet | 6.5 | 0.11 | 12 | 0.064 (0.003-0.304) | 0.158 (0.007-0.747) |
| Arcida | Glycymerididae | Glycymeris | *Glycymeris nummaria* | Violet bittersweet | 7 | 0.11 | 12 | 0.063 (0.003-0.298) | 0.155 (0.007-0.718) |
| Arcida | Glycymerididae | Glycymeris | *Glycymeris ovata* | Black bittersweet | 4.15 | 0.11 | 12 | 0.068 (0.003-0.32) | 0.167 (0.008-0.785) |
| Arcida | Arcidae | Lunarca | *Lunarca ovalis* | Blood ark | 7.6 | 0.45 | 25 | 0.049 (0.002-0.234) | 0.122 (0.006-0.566) |
| Arcida | Arcidae | Tegillarca | *Tegillarca granosa* | Blood cockle | 9 | 1.055 | 28 | 0.118 (0.005-0.571) | 0.289 (0.012-1.424) |
| Cardiida | Cardiidae | Acanthocardia | *Acanthocardia spinosa* | Sand cockle | 8.25 | 0.669 | 26.5 | 0.014 (0.001-0.071) | 0.034 (0.003-0.175) |
| Cardiida | Cardiidae | Acanthocardia | *Acanthocardia tuberculata* | Tuberculate cockle | 9 | 0.669 | 26.5 | 0.006 (0.001-0.016) | 0.015 (0.003-0.044) |
| Cardiida | Psammobiidae | Asaphis | *Asaphis violascens* | Pacific asaphis | 11 | 0.825 | 26.5 | 0.068 (0.004-0.295) | 0.167 (0.011-0.729) |
| Cardiida | Cardiidae | Cerastoderma | *Cerastoderma edule* | Common edible cockle | 5.6 | 0.62 | 10 | 0.101 (0.032-0.243) | 0.258 (0.071-0.653) |
| Cardiida | Cardiidae | Clinocardium | *Clinocardium nuttallii* | Basket cockle | 14 | 0.1525 | 7 | 0.046 (0.003-0.208) | 0.118 (0.008-0.533) |
| Cardiida | Cardiidae | Dallocardia | *Dallocardia muricata* | American yellow cockle | 6 | 0.1525 | 26.5 | 0.022 (0.001-0.11) | 0.055 (0.002-0.277) |
| Cardiida | Donacidae | Donax | *Donax dentifer* | Toothed donax | 3.05 | 0.62 | 26.5 | 0.037 (0.005-0.143) | 0.092 (0.011-0.374) |
| Cardiida | Donacidae | Donax | *Donax obesulus* | Common peruvian donax | 3.9 | 1 | 26.5 | 0.071 (0.007-0.283) | 0.178 (0.018-0.723) |
| Cardiida | Donacidae | Donax | *Donax trunculus* | Truncate donax | 4.4 | 0.7 | 26.5 | 0.035 (0.006-0.114) | 0.089 (0.015-0.298) |
| Cardiida | Psammobiidae | Hiatula | *Hiatula diphos* | Diphos sanguin | 6 | 0.825 | 26.5 | 0.067 (0.017-0.192) | 0.168 (0.041-0.478) |

| | | | | | | | | | |
|---|---|---|---|---|---|---|---|---|---|
| Cardiida | Cardiidae | Hippopus | *Hippopus hippopus* | Bear paw clam | 50 | 0.14 | 26.5 | 0.022 (0-0.124) | 0.056 (0.001-0.311) |
| Cardiida | Cardiidae | Laevicardium | *Laevicardium crassum* | Norwegian egg cockle | 7.5 | 0.1525 | 26.5 | 0.021 (0.001-0.111) | 0.052 (0.002-0.268) |
| Cardiida | Semelidae | Scrobicularia | *Scrobicularia plana* | Peppery furrow | 6.5 | 0.36 | 8 | 0.096 (0.006-0.445) | 0.242 (0.013-1.126) |
| Cardiida | Semelidae | Semele | *Semele solida* | Chilean semele | 3.035 | 0.297 | 12 | 0.068 (0.004-0.332) | 0.171 (0.01-0.84) |
| Cardiida | Solecurtidae | Tagelus | *Tagelus dombeii* | Dombey's tagelus | 7.8 | 0.4 | 26.5 | 0.035 (0.002-0.165) | 0.088 (0.005-0.414) |
| Cardiida | Cardiidae | Tridacna | *Tridacna crocea* | Crocus giant clam | 15 | 0.165 | 26.5 | 0.02 (0.001-0.111) | 0.051 (0.002-0.288) |
| Cardiida | Cardiidae | Tridacna | *Tridacna derasa* | Smooth giant clam | 60 | 0.109 | 26.5 | 0.025 (0-0.171) | 0.062 (0-0.404) |
| Cardiida | Cardiidae | Tridacna | *Tridacna gigas* | Giant clam | 137 | 0.07 | 26.5 | 5.402 (0-0.994) | 12.555 (0-2.478) |
| Cardiida | Cardiidae | Tridacna | *Tridacna maxima* | Elongate giant clam | 41.7 | 0.235 | 26.5 | 0.022 (0-0.136) | 0.055 (0.001-0.336) |
| Cardiida | Cardiidae | Tridacna | *Tridacna squamosa* | Fluted giant clam | 45 | 0.165 | 26.5 | 0.022 (0-0.128) | 0.055 (0.001-0.325) |
| Limida | Limidae | Lima | *Lima lima* | Spiny file shell | 9 | 0.43 | 17 | 0.067 (0.004-0.301) | 0.166 (0.01-0.747) |
| Limida | Limidae | Limaria | *Limaria hians* | Limaria hians | 0.605 | 0.43 | 17 | 0.075 (0.004-0.373) | 0.188 (0.011-0.89) |
| Myida | Myidae | Mya | *Mya arenaria* | Sand gaper | 10 | 0.3 | 10 | 0.055 (0.015-0.146) | 0.14 (0.034-0.397) |
| Mytilida | Mytilidae | Choromytilus | *Choromytilus chorus* | Choro mussel | 26 | 0.58 | 12 | 0.119 (0.009-0.531) | 0.296 (0.022-1.281) |
| Mytilida | Mytilidae | Geukensia | *Geukensia demissa* | Atlantic ribbed mussel | 12 | 0.26 | 16 | 0.065 (0.005-0.284) | 0.161 (0.011-0.673) |
| Mytilida | Mytilidae | Lithophaga | *Lithophaga lithophaga* | European date mussel | 5.6 | 0.044 | 16 | 0.052 (0.003-0.255) | 0.131 (0.007-0.622) |
| Mytilida | Mytilidae | Modiolus | *Modiolus barbatus* | Bearded horse mussel | 6.611 | 0.1955 | 16 | 0.062 (0.004-0.275) | 0.152 (0.01-0.698) |
| Mytilida | Mytilidae | Mytella | *Mytella guyanensis* | Guyana swamp mussel | 6 | 0.128 | 18 | 0.052 (0.003-0.232) | 0.129 (0.007-0.56) |
| Mytilida | Mytilidae | Mytella | *Mytella strigata* | Strigate mangrove mussel | 6 | 0.128 | 18 | 0.052 (0.003-0.23) | 0.131 (0.008-0.566) |
| Mytilida | Mytilidae | Mytilus | *Mytilus californianus* | Californian mussel | 25.5 | 0.25 | 15 | 0.034 (0.015-0.061) | 0.088 (0.034-0.169) |
| Mytilida | Mytilidae | Mytilus | *Mytilus chilensis* | Chilean mussel | 18 | 0.25 | 12 | 0.06 (0.032-0.111) | 0.149 (0.081-0.268) |
| Mytilida | Mytilidae | Mytilus | *Mytilus coruscus* | Korean mussel | 10.5 | 0.25 | 15 | 0.053 (0.012-0.143) | 0.132 (0.029-0.351) |
| Mytilida | Mytilidae | Mytilus | *Mytilus edulis* | Blue mussel | 11 | 0.24 | 8 | 0.078 (0.043-0.13) | 0.195 (0.113-0.318) |
| Mytilida | Mytilidae | Mytilus | *Mytilus galloprovincialis* | Mediterranean mussel | 16.5 | 0.26 | 17 | 0.042 (0.026-0.064) | 0.105 (0.061-0.17) |
| Mytilida | Mytilidae | Mytilus | *Mytilus planulatus* | Australian mussel | 6 | 0.25 | 26 | 0.034 (0.006-0.115) | 0.084 (0.014-0.283) |
| Mytilida | Mytilidae | Mytilus | *Mytilus platensis* | River Plata mussel | 9 | 0.38 | 14 | 0.07 (0.016-0.187) | 0.174 (0.038-0.444) |
| Mytilida | Mytilidae | Perna | *Perna canaliculus* | New Zealand mussel | 15 | 0.6 | 26 | 0.064 (0.016-0.18) | 0.156 (0.042-0.43) |
| Mytilida | Mytilidae | Perna | *Perna perna* | South American rock mussel | 17 | 0.13 | 26 | 0.037 (0.003-0.172) | 0.091 (0.008-0.418) |
| Mytilida | Mytilidae | Perna | *Perna viridis* | Green mussel | 16.5 | 1.07 | 26 | 0.122 (0.042-0.287) | 0.299 (0.114-0.657) |
| Ostreida | Pinnidae | Atrina | *Atrina maura* | Maura pen shell | 29.75 | 0.77 | 20 | 0.109 (0.006-0.526) | 0.269 (0.014-1.26) |
| Ostreida | Ostreidae | Crassostrea | *Crassostrea corteziensis* | Cortez oyster | 25 | 1.09 | 27 | 0.142 (0.02-0.517) | 0.347 (0.051-1.17) |
| Ostreida | Ostreidae | Crassostrea | *Crassostrea rhizophorae* | Mangrove cupped oyster | 12 | 2.79 | 27 | 29.998 (0.018-169.971) | 70.492 (0.047-391.851) |
| Ostreida | Ostreidae | Crassostrea | *Crassostrea tulipa* | Gasar cupped oyster | 25 | 0.845 | 27 | 0.09 (0.016-0.289) | 0.22 (0.039-0.691) |
| Ostreida | Ostreidae | Crassostrea | *Crassostrea virginica* | American cupped oyster | 30 | 0.565 | 24 | 0.054 (0.021-0.115) | 0.135 (0.055-0.281) |
| Ostreida | Ostreidae | Magallana | *Magallana bilineata* | Slipper cupped oyster | 20.88 | 1 | 28 | 0.129 (0.021-0.433) | 0.31 (0.055-0.988) |
| Ostreida | Ostreidae | Magallana | *Magallana gigas* | Pacific cupped oyster | 45 | 0.68 | 13 | 0.123 (0.066-0.213) | 0.303 (0.189-0.464) |
| Ostreida | Ostreidae | Magallana | *Magallana sikamea* | Kumamoto oyster | 6 | 0.84 | 20.5 | 0.186 (0.025-0.67) | 0.449 (0.064-1.496) |
| Ostreida | Ostreidae | Ostrea | *Ostrea conchaphila* | Olympia oyster | 9 | 0.7625 | 14.5 | 0.228 (0.013-1.04) | 0.557 (0.033-2.498) |
| Ostreida | Margaritidae | Pinctada | *Pinctada maxima* | Silverlip pearl oyster | 30 | 0.74 | 22 | 0.09 (0.005-0.417) | 0.219 (0.012-1.014) |
| Ostreida | Margaritidae | Pinctada | *Pinctada radiata* | Rayed pearl oyster | 10.05 | 0.414 | 22 | 0.064 (0.004-0.315) | 0.158 (0.009-0.753) |
| Ostreida | Pteriidae | Pteria | *Pteria penguin* | Penguin wing oyster | 30 | 0.69 | 22 | 0.08 (0.004-0.375) | 0.199 (0.01-0.904) |
| Ostreida | Ostreidae | Saccostrea | *Saccostrea cucullata* | Hooded oyster | 20 | 0.7625 | 24 | 0.148 (0.021-0.491) | 0.362 (0.052-1.147) |
| Pectinida | Pectinidae | Aequipecten | *Aequipecten opercularis* | Queen scallop | 11 | 0.645 | 12 | 0.094 (0.008-0.413) | 0.231 (0.02-0.999) |
| Pectinida | Pectinidae | Argopecten | *Argopecten purpuratus* | Peruvian calico scallop | 12 | 1.995 | 21 | 1.42 (0.01-8.773) | 3.459 (0.026-21.509) |

| | | | | | | | | | |
|---|---|---|---|---|---|---|---|---|---|
| Pectinida | Pectinidae | Argopecten | *Argopecten ventricosus* | Pacific calico scallop | 10 | 2.1 | 26 | 1.388 (0.008-8.634) | 3.322 (0.021-21.089) |
| Pectinida | Pectinidae | Chlamys | *Chlamys islandica* | Iceland scallop | 11 | 0.139 | 8 | 0.054 (0.004-0.257) | 0.135 (0.01-0.617) |
| Pectinida | Pectinidae | Flexopecten | *Flexopecten glaber* | Smooth scallop | 8.66 | 0.5035 | 16 | 0.059 (0.005-0.243) | 0.147 (0.013-0.602) |
| Pectinida | Pectinidae | Mizuhopecten | *Mizuhopecten yessoensis* | Yesso scallop | 25 | 0.925 | 18 | 0.04 (0.017-0.081) | 0.101 (0.043-0.205) |
| Pectinida | Pectinidae | Nodipecten | *Nodipecten nodosus* | Lion's paw | 15 | 0.5035 | 16 | 0.048 (0.007-0.181) | 0.118 (0.018-0.421) |
| Pectinida | Pectinidae | Patinopecten | *Patinopecten caurinus* | Weathervane scallop | 28 | 0.413 | 8 | 0.069 (0.006-0.306) | 0.173 (0.014-0.754) |
| Pectinida | Pectinidae | Pecten | *Pecten fumatus* | Southern Australia scallop | 9.3 | 0.5 | 16 | 0.059 (0.005-0.249) | 0.146 (0.013-0.607) |
| Pectinida | Pectinidae | Pecten | *Pecten jacobaeus* | Great Mediterranean scallop | 25 | 0.5 | 20 | 0.041 (0.003-0.178) | 0.102 (0.008-0.426) |
| Pectinida | Pectinidae | Pecten | *Pecten maximus* | Great Atlantic scallop | 17 | 0.475 | 12 | 0.065 (0.006-0.278) | 0.162 (0.015-0.687) |
| Pectinida | Pectinidae | Placopecten | *Placopecten magellanicus* | American sea scallop | 20 | 0.28 | 8 | 0.013 (0.006-0.024) | 0.034 (0.014-0.07) |
| Pectinida | Spondylidae | Spondylus | *Spondylus americanus* | Atlantic thorny oyster | 11.2 | 0.5 | 16 | 0.07 (0.005-0.325) | 0.175 (0.012-0.763) |
| Pectinida | Spondylidae | Spondylus | *Spondylus crassisquama* | Pacific thorny oyster | 8.85 | 0.5 | 16 | 0.074 (0.005-0.341) | 0.184 (0.012-0.822) |
| Pectinida | Pectinidae | Zygochlamys | *Zygochlamys delicatula* | Delicate scallop | 7.8 | 0.187 | 13 | 0.045 (0.003-0.203) | 0.112 (0.008-0.495) |
| Venerida | Arcticidae | Arctica | *Arctica islandica* | Ocean quahog | 13 | 0.0605 | 8 | 0.056 (0.003-0.28) | 0.138 (0.008-0.669) |
| Venerida | Veneridae | Austrovenus | *Austrovenus stutchburyi* | Stutchbury's venus | 4.9 | 0.3115 | 15 | 0.057 (0.005-0.231) | 0.141 (0.011-0.573) |
| Venerida | Veneridae | Callista | *Callista chione* | Smooth callista | 11 | 0.24 | 12 | 0.04 (0.008-0.122) | 0.102 (0.02-0.319) |
| Venerida | Veneridae | Chamelea | *Chamelea gallina* | Striped venus | 5 | 0.415 | 14 | 0.045 (0.009-0.135) | 0.113 (0.022-0.352) |
| Venerida | Veneridae | Cyclina | *Cyclina sinensis* | Oriental cyclina | 5 | 0.421 | 16 | 0.063 (0.005-0.261) | 0.157 (0.013-0.612) |
| Venerida | Veneridae | Dosinia | *Dosinia exoleta* | Mature dosinia | 4.615 | 0.448 | 16 | 0.065 (0.006-0.265) | 0.16 (0.014-0.637) |
| Venerida | Veneridae | Gafrarium | *Gafrarium tumidum* | Tumid venus | 4 | 0.32565 | 16 | 0.054 (0.005-0.213) | 0.133 (0.011-0.52) |
| Venerida | Veneridae | Iliochione | *Iliochione subrugosa* | Semi-rough venus | 4.485 | 0.421 | 16 | 0.064 (0.006-0.265) | 0.159 (0.014-0.627) |
| Venerida | Veneridae | Leukoma | *Leukoma staminea* | Pacific littleneck clam | 7.5 | 0.21105 | 8 | 0.07 (0.006-0.295) | 0.174 (0.014-0.718) |
| Venerida | Veneridae | Leukoma | *Leukoma thaca* | Taca clam | 8 | 0.174 | 19 | 0.036 (0.003-0.154) | 0.089 (0.007-0.372) |
| Venerida | Mactridae | Lutraria | *Lutraria magna* | Oblong otter shell | 5.115 | 0.1975 | 11 | 0.043 (0.003-0.198) | 0.107 (0.007-0.493) |
| Venerida | Mactridae | Mactra | *Mactra quadrangularis* | Globose clam | 6.05 | 0.15 | 11 | 0.04 (0.003-0.188) | 0.099 (0.006-0.461) |
| Venerida | Mactridae | Mactromeris | *Mactromeris polynyma* | Stimpson's surf clam | 16 | 0.0801 | 4 | 0.05 (0.003-0.239) | 0.124 (0.007-0.591) |
| Venerida | Veneridae | Mercenaria | *Mercenaria campechiensis* | Southern hardshell clam | 13 | 0.64 | 17 | 0.149 (0.017-0.47) | 0.363 (0.041-1.081) |
| Venerida | Veneridae | Mercenaria | *Mercenaria mercenaria* | Northern quahog(=Hard clam) | 13 | 0.3257 | 17 | 0.102 (0.037-0.225) | 0.248 (0.106-0.498) |
| Venerida | Veneridae | Meretrix | *Meretrix lusoria* | Japanese hard clam | 5 | 1.26 | 17 | 0.34 (0.011-1.774) | 0.828 (0.028-4.243) |
| Venerida | Mesodesmatidae | Mesodesma | *Mesodesma donacium* | Macha clam | 7 | 1.13 | 17 | 0.232 (0.009-1.229) | 0.569 (0.022-2.981) |
| Venerida | Mesodesmatidae | Paphies | *Paphies australis* | Pipi wedge clam | 8 | 0.16 | 20 | 0.036 (0.002-0.178) | 0.089 (0.005-0.432) |
| Venerida | Veneridae | Polititapes | *Polititapes aureus* | Golden carpet shell | 6 | 0.421 | 12 | 0.08 (0.007-0.348) | 0.198 (0.017-0.816) |
| Venerida | Veneridae | Polititapes | *Polititapes rhomboides* | Banded carpet shell | 6 | 0.421 | 12 | 0.08 (0.007-0.332) | 0.197 (0.018-0.794) |
| Venerida | Veneridae | Ruditapes | *Ruditapes decussatus* | Grooved carpet shell | 8 | 0.48 | 17 | 0.033 (0.011-0.087) | 0.082 (0.028-0.198) |
| Venerida | Veneridae | Ruditapes | *Ruditapes philippinarum* | Japanese carpet shell | 8 | 0.52 | 18 | 0.031 (0.015-0.058) | 0.075 (0.042-0.122) |
| Venerida | Veneridae | Saxidomus | *Saxidomus gigantea* | Butter clam | 13 | 0.421 | 7 | 0.103 (0.007-0.439) | 0.255 (0.019-1.057) |
| Venerida | Mactridae | Spisula | *Spisula murchisoni* | Large trough shell | 5 | 0.3307 | 12.5 | 0.018 (0.003-0.079) | 0.043 (0.007-0.192) |
| Venerida | Mactridae | Spisula | *Spisula sachalinensis* | Imperial surf clam | 5 | 0.3307 | 18 | 0.013 (0.002-0.063) | 0.032 (0.005-0.149) |
| Venerida | Mactridae | Spisula | *Spisula solida* | Solid surf clam | 5 | 0.3307 | 11 | 0.01 (0.004-0.022) | 0.024 (0.01-0.051) |
| Venerida | Mactridae | Spisula | *Spisula subtruncata* | Subtruncate surf clam | 3 | 0.67 | 12.5 | 0.033 (0.004-0.152) | 0.079 (0.011-0.36) |

| | | | | | | | | |
|---|---|---|---|---|---|---|---|---|
| Venerida | Veneridae | Tawera | *Tawera elliptica* | Gay's little venus | 2.803 | 0.288 | 12 | 0.066 (0.006-0.284) | 0.163 (0.014-0.682) |
| Venerida | Veneridae | Tivela | *Tivela mactroides* | Triangular tivela | 3.4 | 0.15 | 27 | 0.027 (0.001-0.132) | 0.068 (0.003-0.32) |
| Venerida | Mactridae | Tresus | *Tresus nuttallii* | Pacific horse clam | 25.4 | 0.153 | 11 | 0.035 (0.002-0.168) | 0.087 (0.004-0.422) |
| Venerida | Veneridae | Venerupis | *Venerupis corrugata* | Corrugated venus | 7.7 | 0.226 | 13 | 0.053 (0.005-0.225) | 0.133 (0.011-0.535) |
| Venerida | Veneridae | Venus | *Venus casina* | Chamber venus | 5 | 0.26 | 19 | 0.043 (0.004-0.183) | 0.106 (0.008-0.442) |
| Venerida | Veneridae | Venus | *Venus verrucosa* | Warty venus | 7 | 0.26 | 19 | 0.042 (0.003-0.176) | 0.103 (0.009-0.434) |

### Supplemental Figures

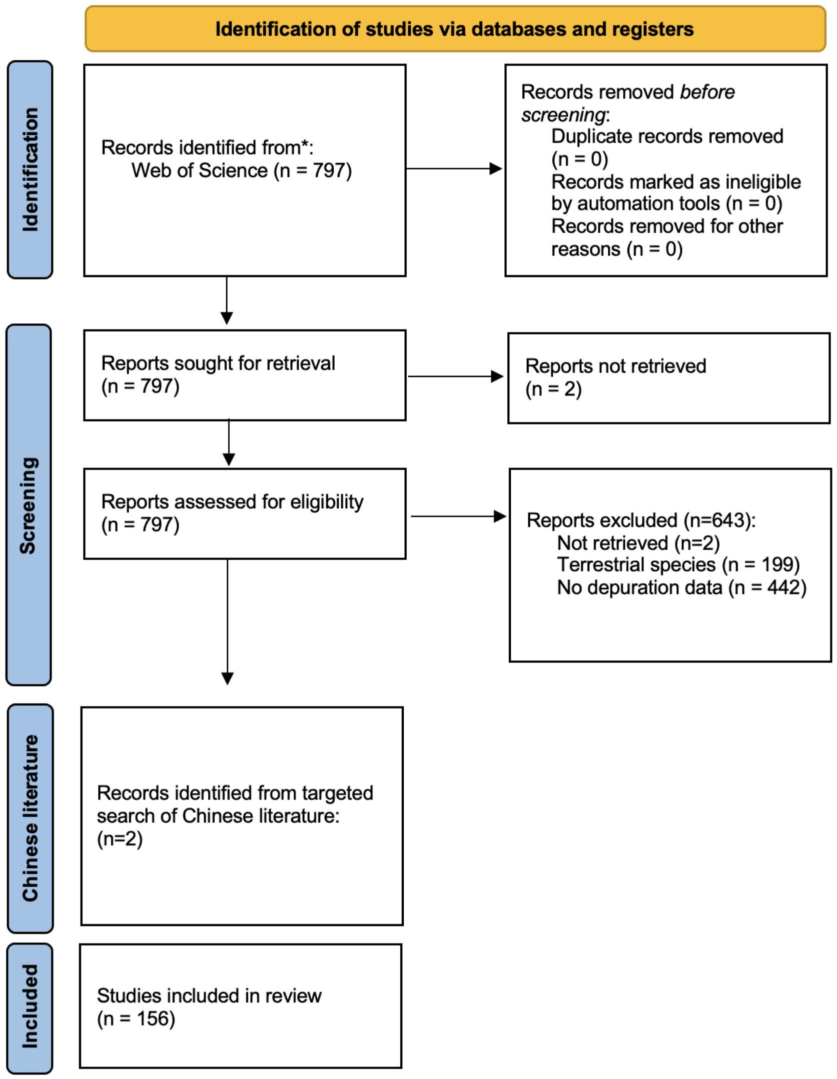


**Figure S1.** PRISMA 2020 flow diagram detailing the literature review inclusion criteria.

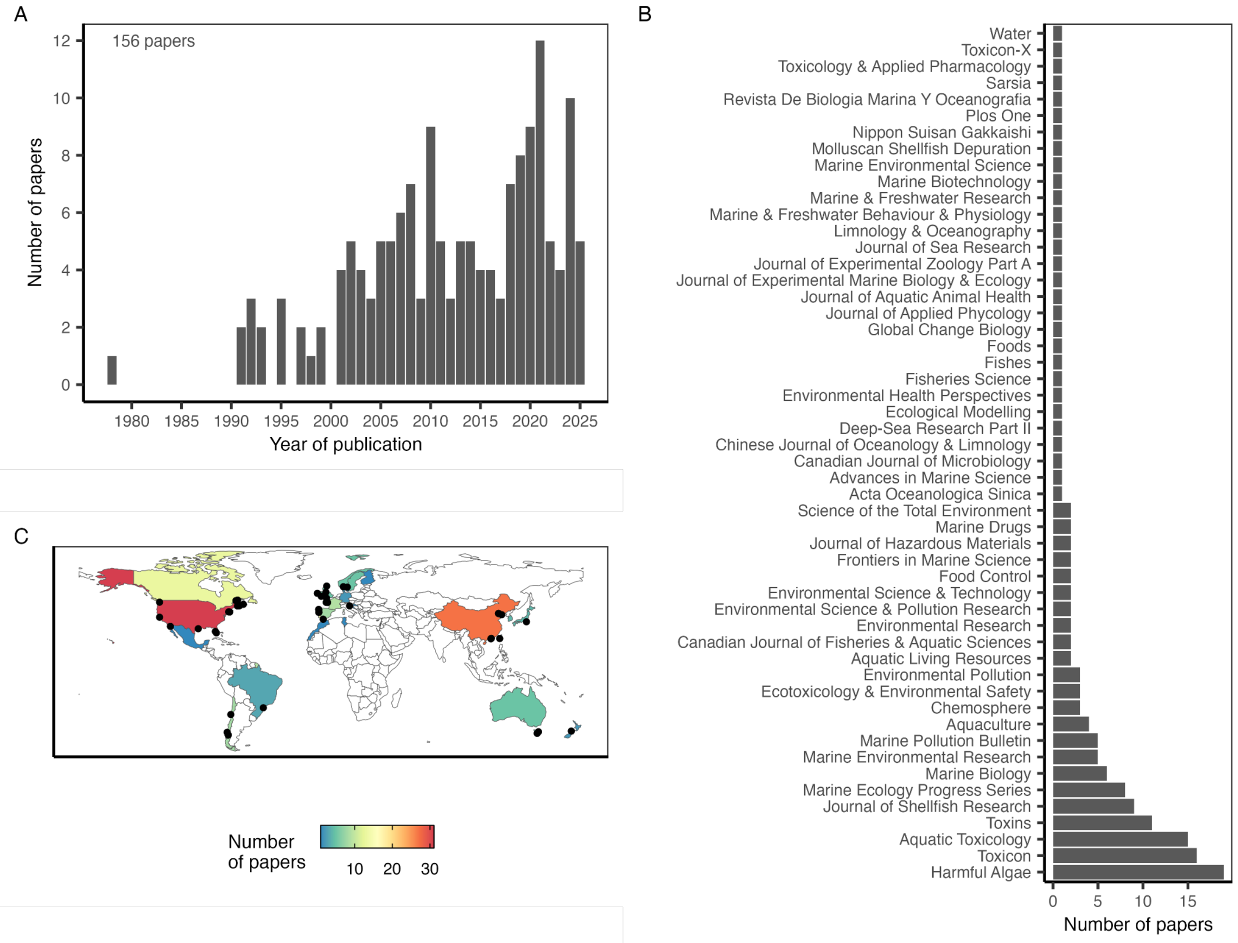


**Figure S2.** The number of papers measuring marine biotoxin depuration rates **(A)** over time, **(B)** by journal; and **(C)** by country of the lead author's primary affiliation. In **(C)**, points mark locations where depuration rates have been quantified from field monitoring data.

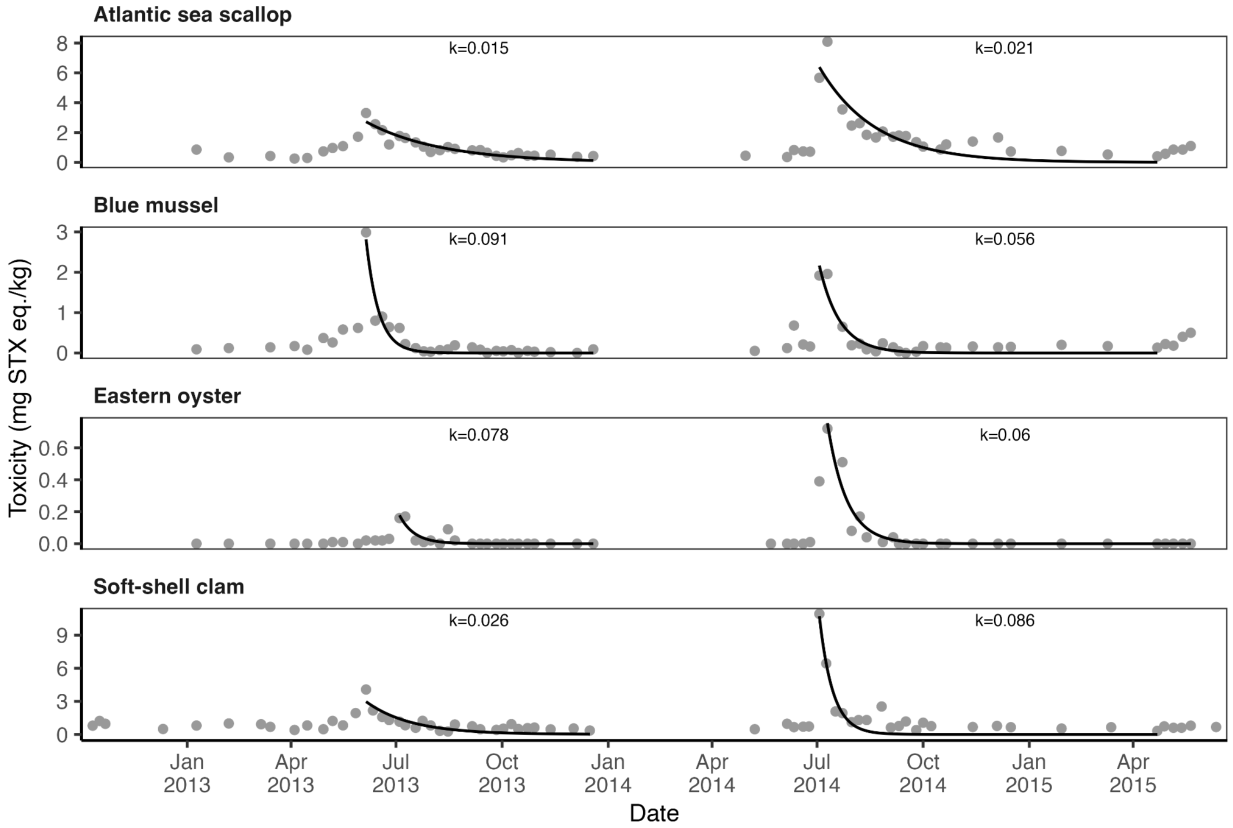


**Figure S3.** An illustration of the methods for estimating paralytic shellfish toxin (PST) depuration rates from biotoxin monitoring data on four shellfish species from Deadmans Harbour, New Brunswick, Canada. Data are from (Rourke et al., 2021). The black line shows the fit of a one-compartment depuration model to the depuration phase of annual toxin events; the exponential decay constant (k) associated with each curve is printed above the curve.

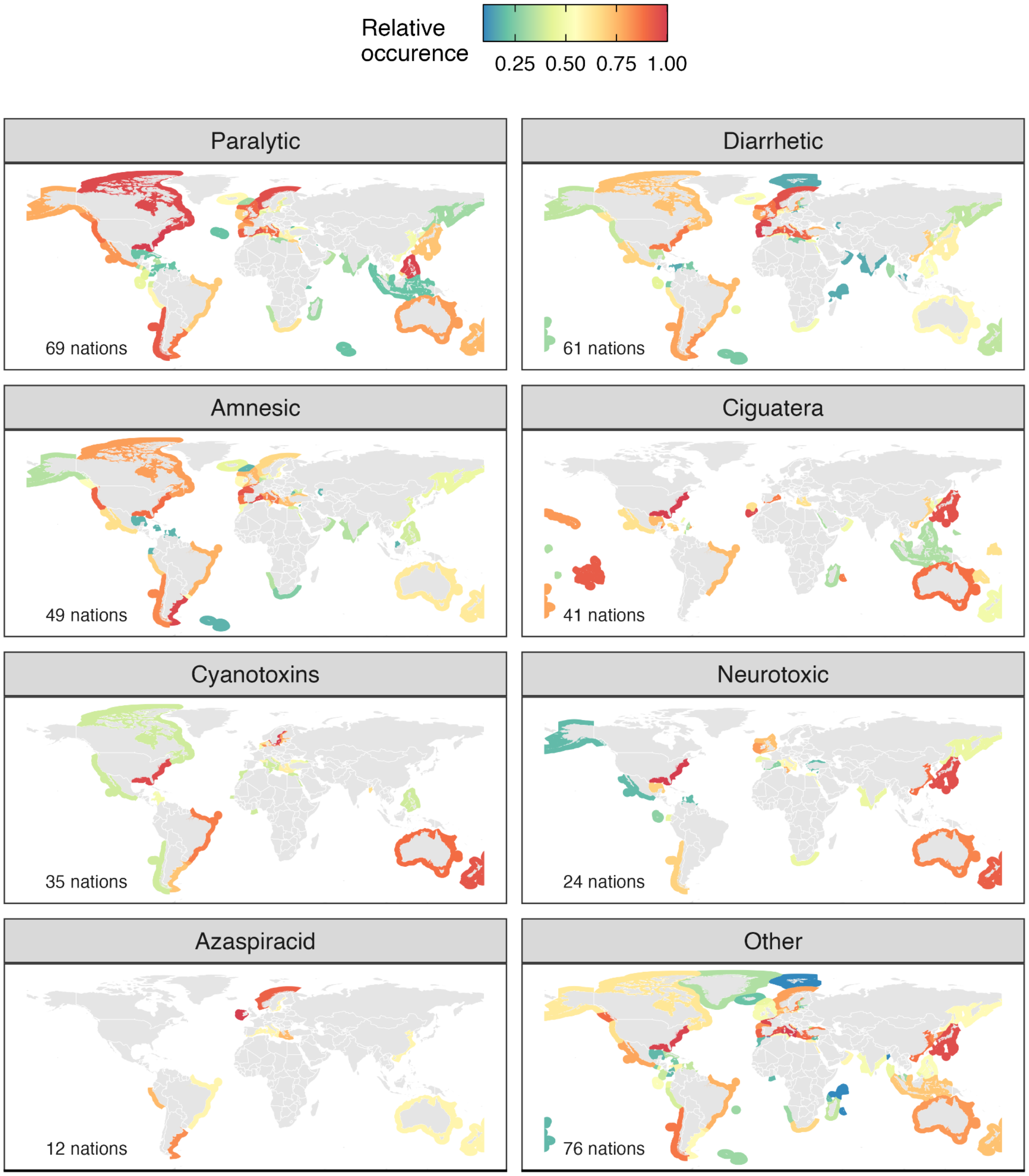


**Figure S4.** Exclusive economic zones (EEZs) exposed to harmful algae and their associated public health risks. Shaded EEZs have HAB species present based on OBIS. Syndromes are sorted by the number of EEZs with observations of HAB species; the number of EEZs impacted (out of 257) is printed in the bottom right.

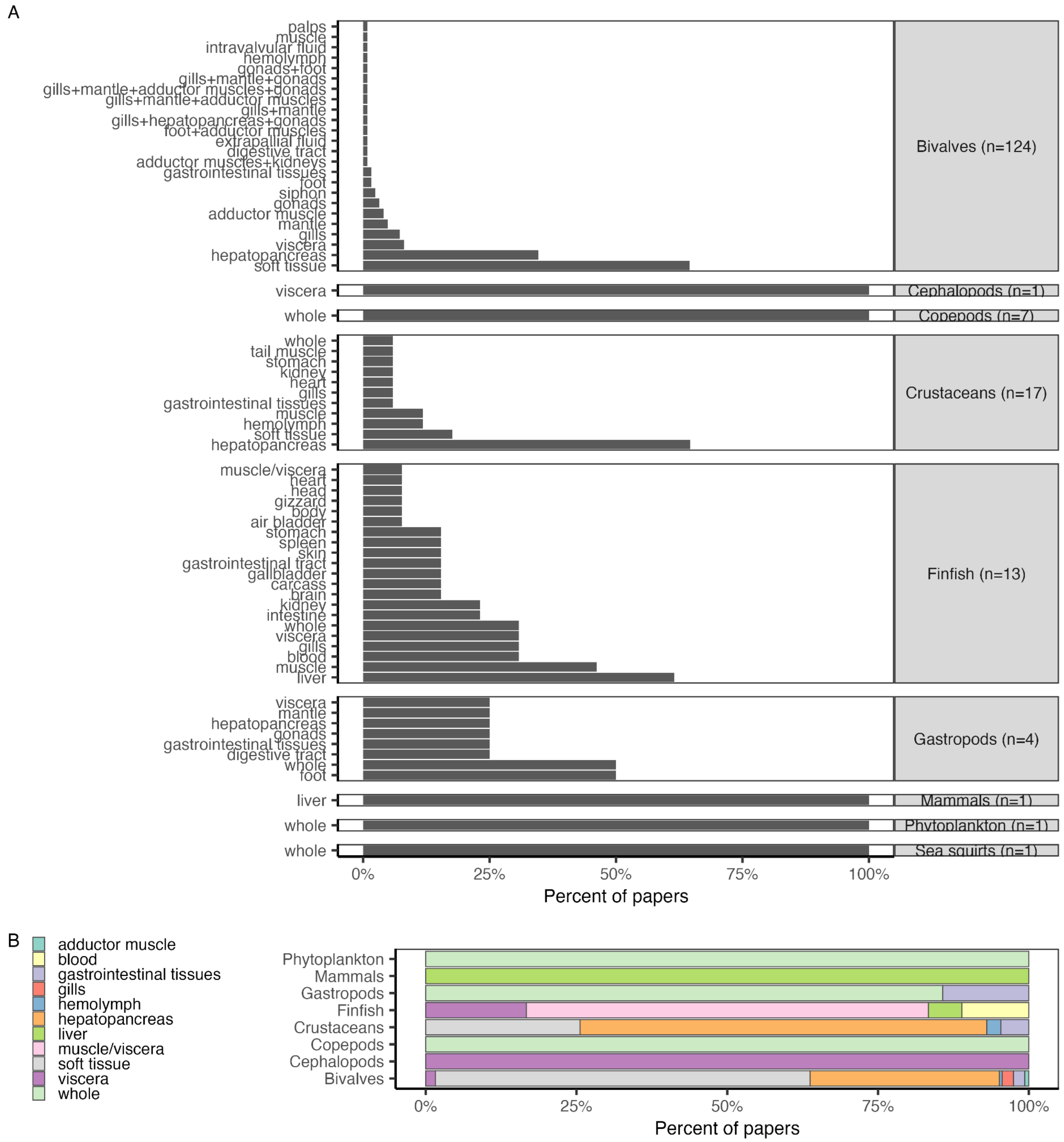


**Figure S5.** Panel **A** shows the frequency with which different tissues are evaluated for biotoxin depuration rates by taxonomic class. The x-axis represents the percent of papers on species within each class evaluating depuration rates for each tissue. The number of papers on species within each class is printed in the class label. Tissues are ordered in increasing frequency (bottom to top). Panel **B** shows the frequency with which different tissues are evaluated for biotoxin depuration rates by taxonomic class in papers in which only a single tissue is evaluated.

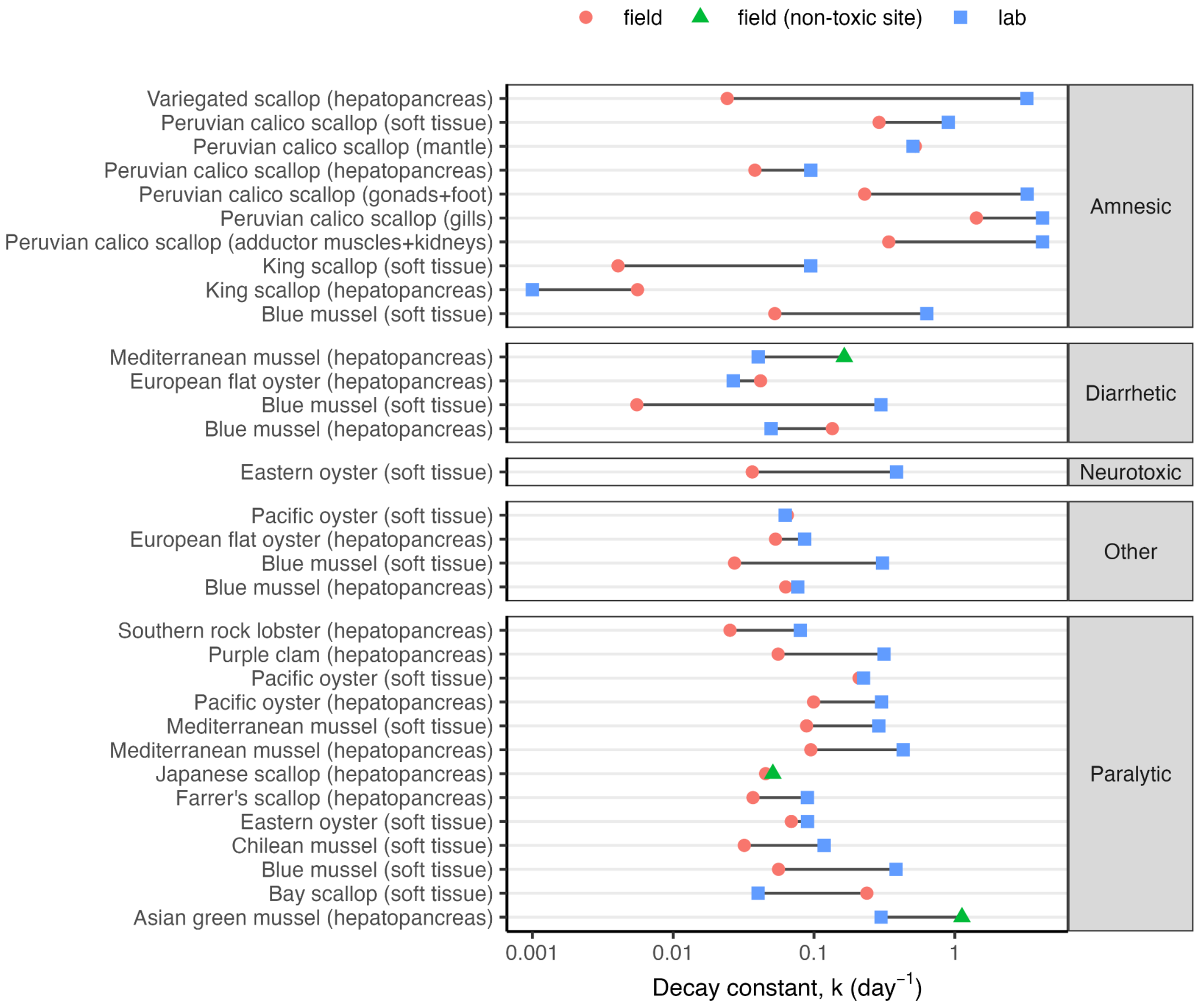


1278

1279 **Figure S6.** A comparison of average depuration rates in the field versus the lab by species,
1280 tissue, and toxin. In general, depuration is faster in the lab than in the field.

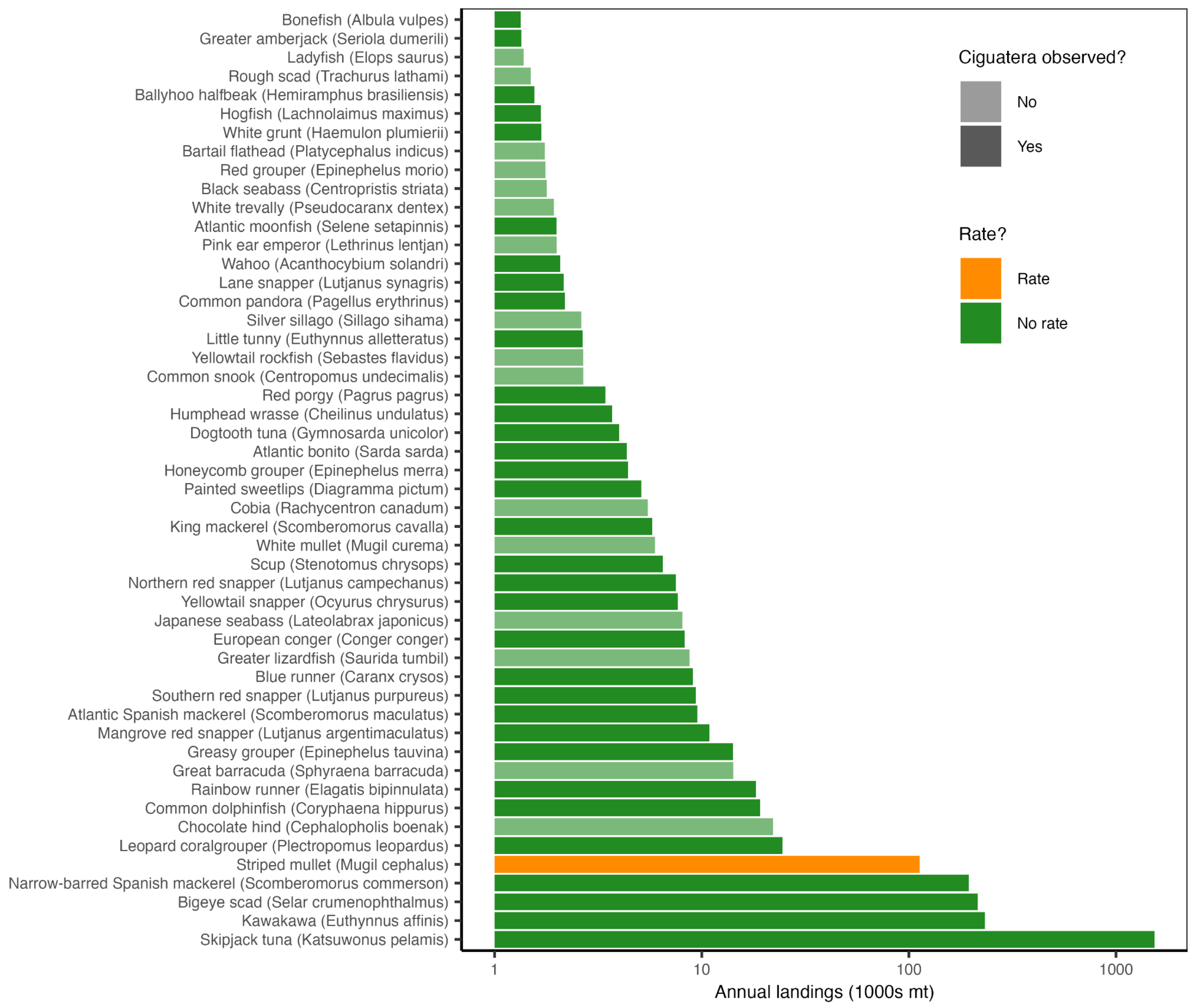


**Figure S7.** The top-50 marine finfish vulnerable to ciguatera most frequently landed by fisheries from 2014-2023 from countries with known ciguatera occurrences. Marine finfish vulnerable to ciguatera were identified as those with reported ciguatera occurrences (known) and large (>25 cm max length), reef-associated, non-herbivorous marine finfish speculated to be vulnerable to ciguatera. Ciguatera depuration rates have not been studied for any of these species.

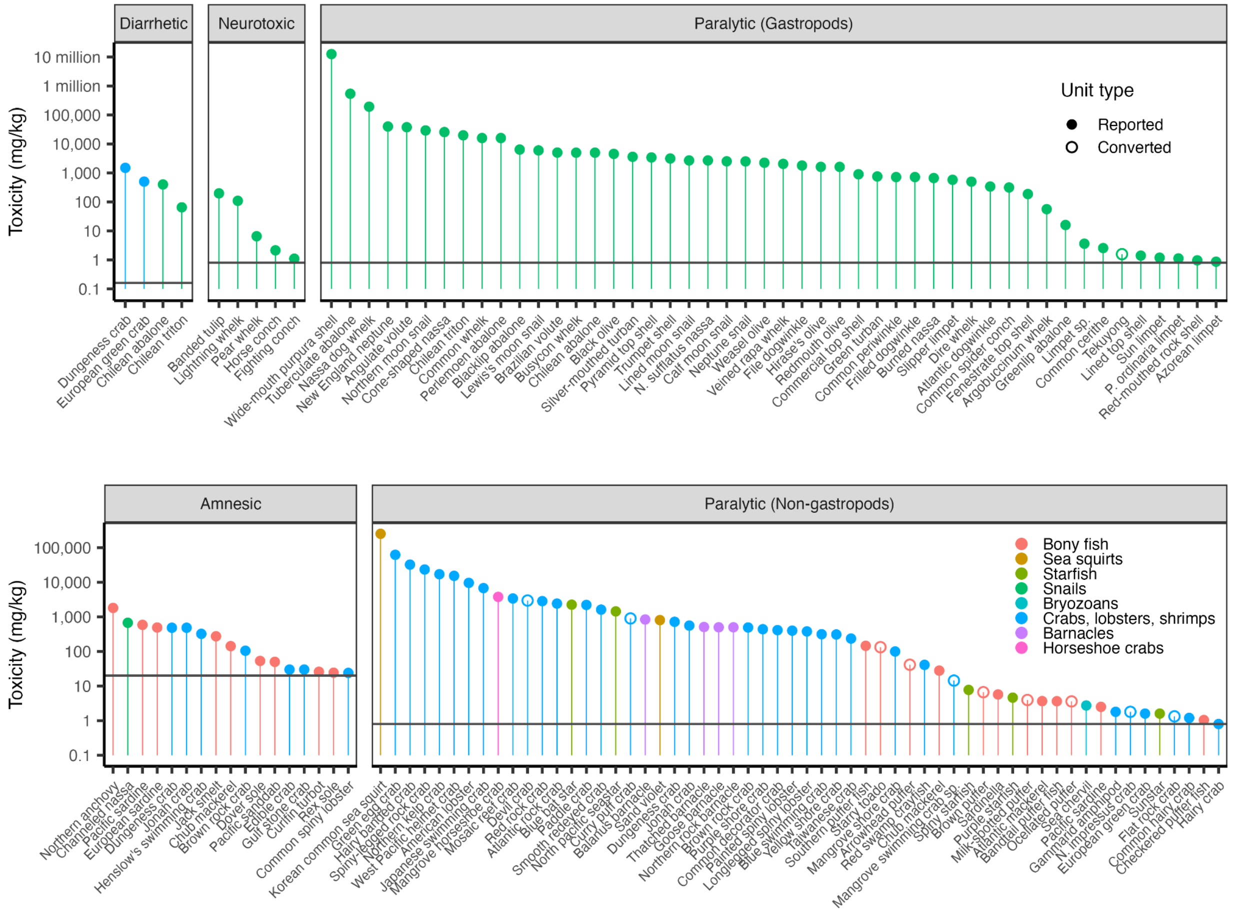


**Figure S8.** Maximum biotoxin toxicities observed in non-bivalve species reported in select review papers (Costa et al., 2017; Deeds et al., 2008; Lefebvre and Robertson, 2010) relative to the most common international action thresholds (Langlois and Morton, 2018), which are indicated by the solid horizontal lines. Note that toxicities in units of mg/kg, µg/g, and ppm are all numerically equivalent (i.e., the y-axis can be directly interpreted as µg/g or ppm). When necessary, we converted PST mouse units (MU) to µg assuming that 1 MU equals 0.18 µg STX equivalents (Arnich and Thébault, 2018); converted toxicities are shown by open circles.

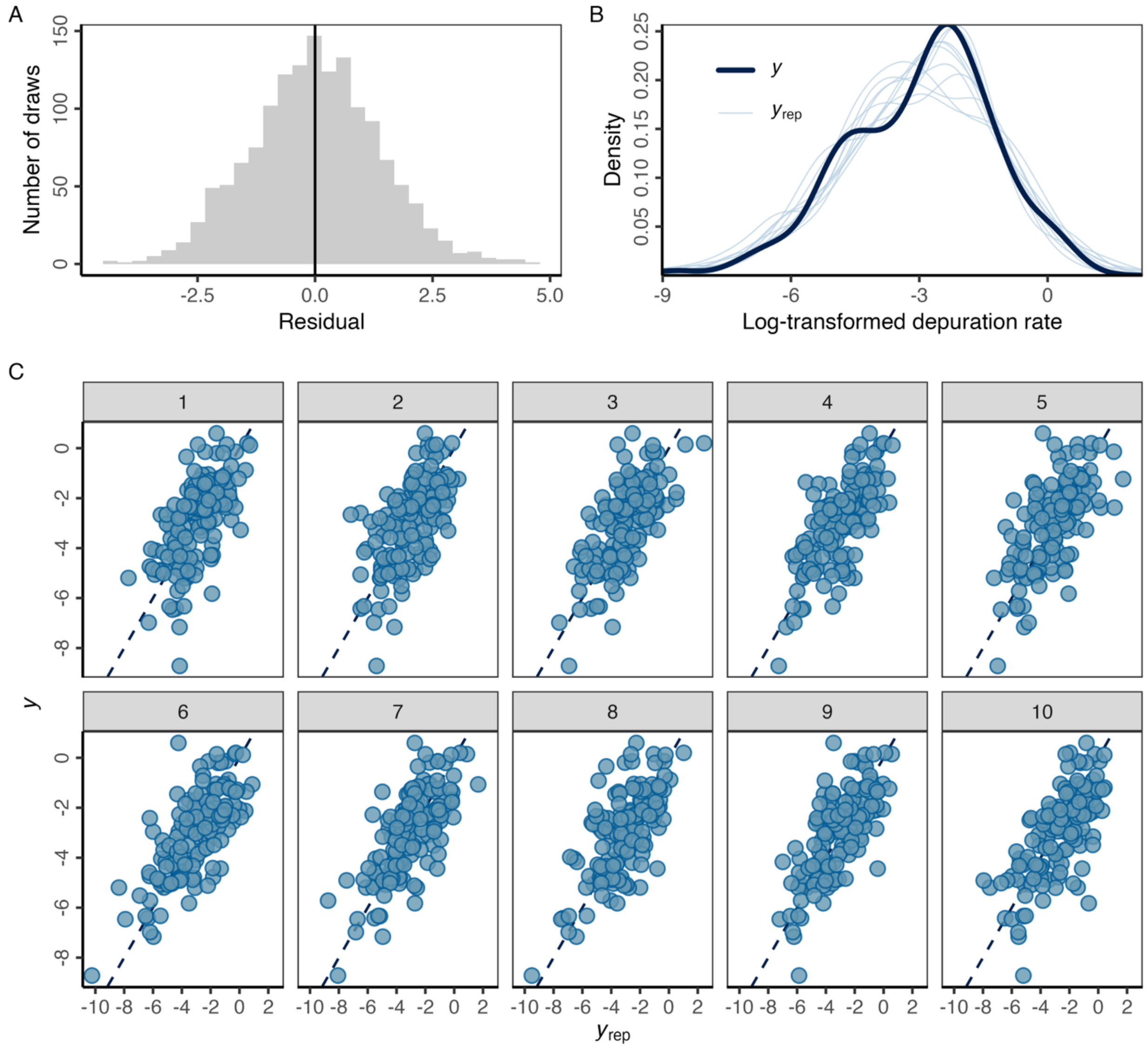


**Figure S9.** Model diagnostic plots for the best performing Bayesian regression model showing **(A)** the distribution of the residuals in 10 posterior draws; **(B)** distribution of the response variable in the data (*y*) versus 10 posterior draws ($y_{rep}$); and **(C)** correlation between the response variable in the data (*y*) versus 10 posterior draws ($y_{rep}$). In **(C)**, the diagonal line represents the one-to-one line and each facet represents a set of posterior draws.

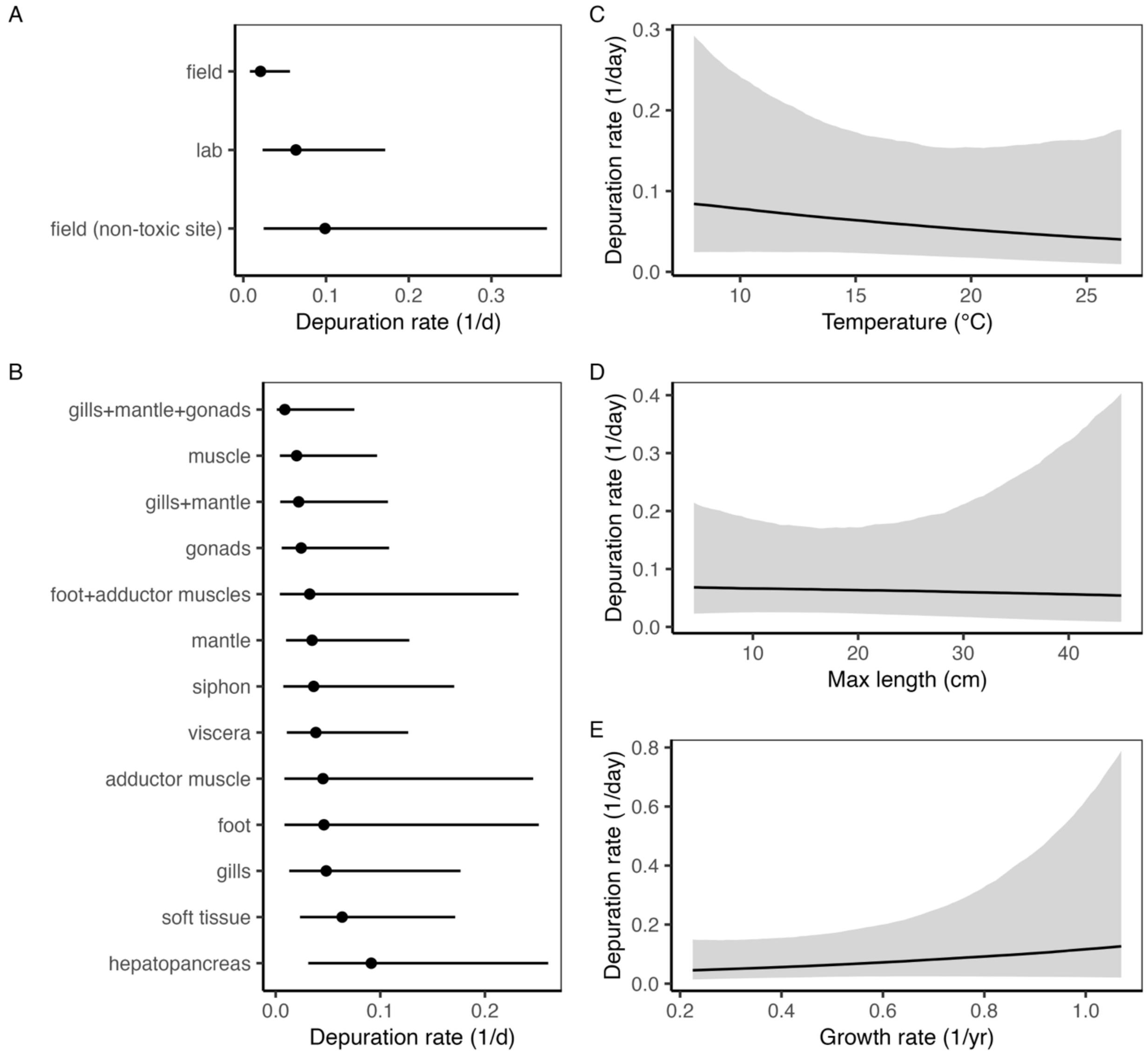


**Figure S10.** The conditional effects of the **(A)** location; **(B)** tissue, (**C)** preferred temperature, **(D)** maximum length, and **(E)** von Bertalanffy growth rate fixed effects variables included in the best performing Bayesian regression model. Conditional effects illustrate the depuration rate expected for each fixed effect value when other fixed and random effects are held at their average. In **(A)** and **(B)**, points represent the median and lines represent the 95% credible interval. In **(C-D)**, lines represent the median and sharing indicates the 95% credible interval.
